# Population size mediates the contribution of high-rate and large-benefit mutations to parallel evolution

**DOI:** 10.1101/2021.02.02.429281

**Authors:** Martijn F. Schenk, Mark P. Zwart, Sungmin Hwang, Philip Ruelens, Edouard Severing, Joachim Krug, J. Arjan G.M. de Visser

**Author notes:** equal contribution (Martijn F. Schenk, Mark P. Zwart).

## Abstract

Both mutations with large benefits and mutations occurring at high rates may cause parallel evolution, but their contribution is expected to depend on population size. We show that small and large bacterial populations adapt to a novel antibiotic using similar numbers, but different types of mutations. Small populations repeatedly substitute similar high-rate structural variants, including the deletion of a nonfunctional β-lactamase, and evolve modest resistance levels. Hundred-fold larger populations more frequently use the same low-rate, large-benefit point mutations, including those activating the β-lactamase, and reach 50-fold higher resistance levels. Our results demonstrate a key role of clonal interference in mediating the contribution of high-rate and large-benefit mutations in populations of different size, facilitated by a tradeoff between rates and fitness effects of different mutation classes.

---

Public health threats from rapidly evolving pathogens, together with observations of convergent and parallel evolution, have stimulated recent efforts to explore the predictability of evolutionary processes [1–5]. Yet, even our understanding of the contribution of fundamental factors such as mutation and selection to parallel evolution is incomplete [6–8]. Mutations occur in various forms and rates and have diverse fitness effects, which make their contribution dependent on population size [9–11]. In sufficiently small populations, high-rate and large-benefit mutations are predicted to impact adaptation similarly [11, 12], while in large populations selection dominates mutation choices, because clonal interference filters out large-effect mutations even when they have low rates [13–15] (Figs. 1 and S1). To what extent high-rate mutations shape adaptation also in large populations is the topic of current debate, largely based on theoretical arguments and observations of transition bias among point mutations [16–20]. Moreover, little is known about the consequences of high-rate and large-benefit mutations for longer-term adaptation.

**Fig. 1.**
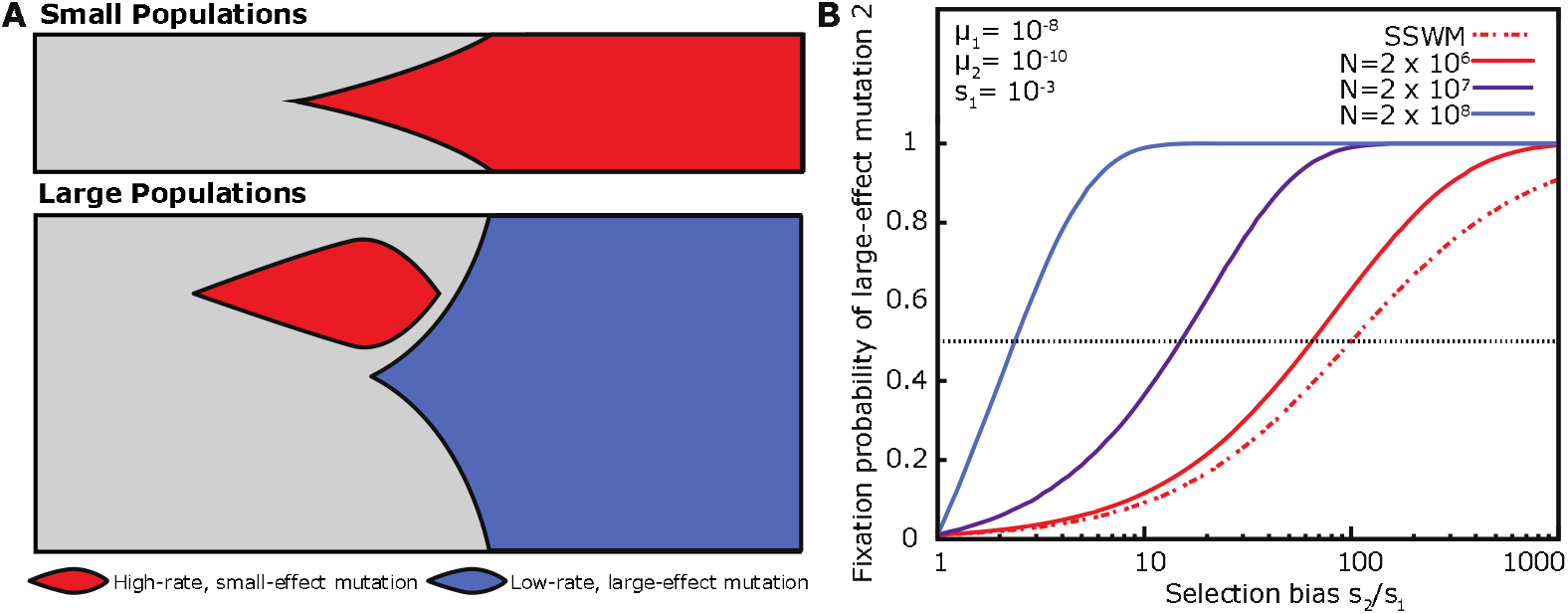
Expected relative impact of high-rate and large-benefit mutations in asexual populations of different size. (**A**) Clonal interference in large populations filters out large-effect beneficial mutations (blue), while in the absence of the latter mutations in small populations high-rate mutations (red) are more likely to dominate. The separation of high-rate and large-benefit mutations in populations of different size is facilitated when mutation rates and effects correlate negatively. (**B**) Effect of population size on the relative fixation probability of two mutations, one (mutation 2) with a low rate (*μ*_2_ = 10^−10^) and up to 1,000-fold larger selective benefit and another (mutation 1) with 100-fold higher rate, but smaller benefit [13]. While in small populations lacking clonal interference (SSWM conditions, Strong Selection Weak Mutation), the relative benefit of mutation 2 should be equal to the inverse of its relative mutation rate to have equal fixation probability as high-rate mutation 1, in populations of the size of our large bacterial populations (2 × 10^8^), a two-fold benefit is sufficient.

We examined the effect of population size on the type, repeatability and adaptive consequences of selected mutations in bacterial populations adapting to a novel antibiotic. Seventy-two small (*N*_e_ ~2 × 10^6^) and 24 large populations (*N*_e_ ~2 × 10^8^) of an *Escherichia coli* strain harboring a multicopy nonconjugative plasmid expressing TEM-1 β-lactamase, evolved via serial transfer in Luria broth containing cefotaxime (CTX) (and tetracycline to avoid plasmid loss). The β-lactamase has very low activity against CTX, but can be activated by point mutations [21]. To maximize selection for resistance, CTX concentrations were increased by a factor of 2^0.25^ whenever the optical density of a population before transfer had risen above 75% of that in the absence of antibiotic, leading to a 4.6-fold higher geometric mean CTX concentration in large than in small populations (Fig. S2). Sixteen large control populations evolved without antibiotics or with only tetracycline (Table S2). After 50 transfers (~500 generations), a random clone was isolated from each population to determine the extent of adaptation. Large populations showed markedly higher resistance levels than small populations (on average 12.8 versus 7.2 doublings of the minimal inhibitory concentration (MIC) of CTX, respectively, *P*<0.001; Fig. 2A).

**Fig. 2.**
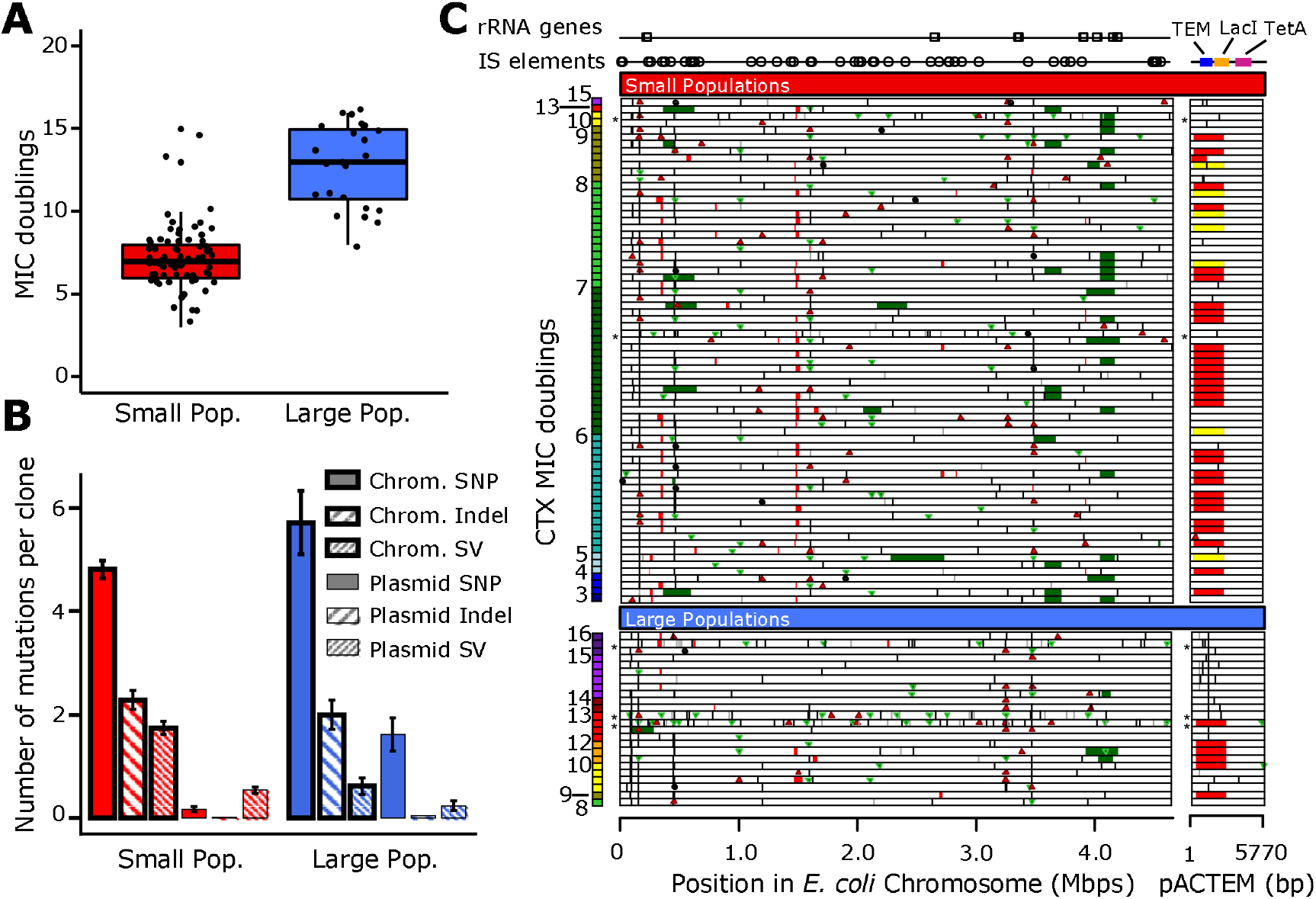
Phenotypic and genetic changes. (**A**) Box plots of the CTX MIC doublings of evolved clones from small (red) and large populations (blue) relative to the ancestral strain. (**B**) Number of mutations per category (means ± SEM), genomic element and population size, for the 91 non-mutator clones. (**C**) Position of mutations in chromosome and plasmid (different scales) for small (top) and large populations (bottom) ranked by their MIC value. Black vertical lines are single nucleotide polymorphisms (SNPs), green and red triangles are small (<1 kbp) insertions and deletions, respectively (indels), black circles are IS-element insertions, red and green bars are large (>1 kbp) deletions and duplications, respectively (SVs, structural variants); yellow bars in the plasmid indicate heteroplasmic deletions; * indicate mutator genotypes; positions of IS-elements and rRNA operons are indicated above.

Resequencing of the ancestral strains and 112 evolved clones revealed 1,190 mutations (Fig. 2B-C, Figs. S5-11 and Tables S4-5). These include 706 single-nucleotide polymorphisms (SNPs), 275 indels (<1kbp) and insertion-sequence (IS) element transpositions, 160 large deletions (>1kbp), 49 large duplications (>1kbp) and four 304-bp inversions). Two clones from the small and three from the large CTX-treated populations were identified as mutators (Fig. S6) [13]. Non-mutator CTX-treated clones had on average 9.5 mutations in small and 10.2 mutations in large populations (*P*=0.405); clones from control populations had fewer mutations (*P*<0.0001, 5.9 in the no-antibiotic and 6.0 in the tetracycline-only populations, Fig. S8). Small populations showed fewer SNPs (*P*<0.0001), particularly in their plasmid, while large populations had fewer structural variants (SVs, i.e. deletions and duplications >1 kbp; *P*<0.0001) in both chromosome and plasmid (Fig. 2B). Of the 503 SNPs observed in CTX-treated non-mutators, 14 were synonymous and 30 intergenic. The normalized ratio of nonsynonymous to synonymous substitutions per site (dN/dS: 11.9 in small, 28.8 in large populations, Table S6) confirmed a dominant role for selection, with 92% and 97% of the nonsynonymous substitutions expected to be beneficial in small and large populations, respectively.

To determine mutational repeatability, the average pairwise similarity of genotypes was calculated for non-mutator clones (Fig. 3A) [13]. Consistent with previous findings [5, 26], mutational repeatability was higher at the gene than at the nucleotide level (34% and 11% shared mutations, respectively). Repeatability was also higher in large than in small populations at the gene (*P*<0.0001) and nucleotide level (*P*=0.032; Fig. 3A). However, SNPs and SVs contribute to this pattern in opposite ways: small populations consistently shared fewer SNPs but more SVs than large populations, both at the nucleotide (Fig. 3B) and gene level (Fig. S14), as well as in chromosome and plasmid (*P*<0.001 in all cases; Tables S7 and S8). What caused this greater repeatability of SNPs in large and SVs in small populations? We hypothesized that a tradeoff between rates and fitness effects of SNPs and SVs underlies this pattern. If SNPs have both lower rates and larger benefits than SVs, clonal interference and stronger purifying selection would more often prevent high-rate SVs from fixing in large than in small populations.

**Fig. 3.**
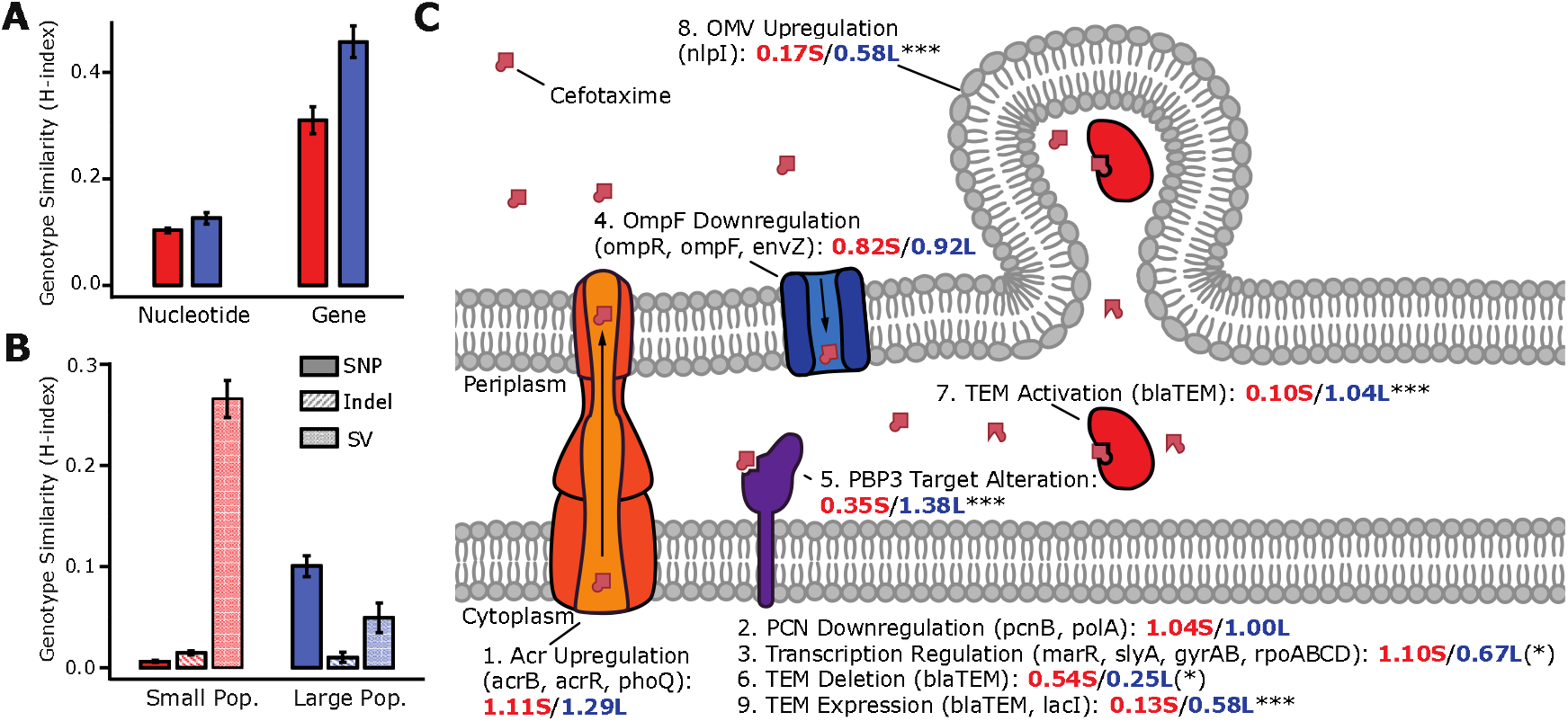
Repeatability and common targets of genomic changes. (**A**) Pairwise mutation similarity (means ± SEM) at the nucleotide and gene level for non-mutator clones, based on H-index [18]. (**B**) Pairwise nucleotide-level mutation similarity per mutation class in chromosome and plasmid (means ± SEM). (**C**) Functional targets with > 20 mutations [13]. Names of genes involved are given between brackets together with the average number of mutations per clone in small (red) and large (blue) populations. Asterisks show result from χ^2^ test of difference in mutation frequency in small and large populations: (*) *P*<0.10, *** *P*<0.001.

Since small and large populations experienced different CTX concentrations (Fig. S2), we first asked whether this had an effect on the choice of SVs and SNPs. Regression analysis showed no effect of variation in CTX concentration on the fraction of SVs for small and large populations separately (*P*≥0.34), only for the combined populations (*P*<0.01, Fig. S15), indicating that differences in mutation supplies rather than CTX concentrations affected the choice of mutations.

To examine whether the adaptive targets were different for small and large populations, we grouped all genes with ≥ five SNPs or indels across all 96 populations into nine functional targets with ≥ 20 mutations (Table S13) [13], which covered 57% of all mutations in these populations. The targets included known β-lactam resistance targets, but also unexpectedly the deletion of *bla*_TEM1_ and its repressor *lacI* from the plasmid (Fig. 3C). All nine targets were affected in small and large populations, albeit in subtly different ways: large populations more often activated the β-lactamase, altered CTX target PBP3 and increased the production of outer-membrane vesicles, while small populations tended to more frequently delete *bla*_TEM1_ and alter transcription regulation. Moreover, as for the total set of mutations (Fig. 2B), also these shared targets were affected more often by SNPs in large populations and by SVs in small populations (*P*<0.0001, Table S13). Thus, small and large populations adapted via similar resistance mechanisms, but differed in the frequency and types of mutations used.

To test our tradeoff hypothesis, we first analyzed the temporal dynamics of mutations based on the metagenomes of five small and five large populations analyzed at 100-generation intervals. The inferred Muller plots (Fig. 4A) show stronger clonal interference in these large populations, where the majority genotype detected at the initial time point never fixes, while it fixes in all small populations (Fisher’s *P*=0.004). SVs are detected earlier than SNPs (*P*=0.042, Fig. 4B), consistent with their expected higher rate [22], but in large populations fewer fix and they do so later than SNPs (*P*=0.015, Fig. 4C), consistent with smaller fitness effects.

**Fig. 4.**
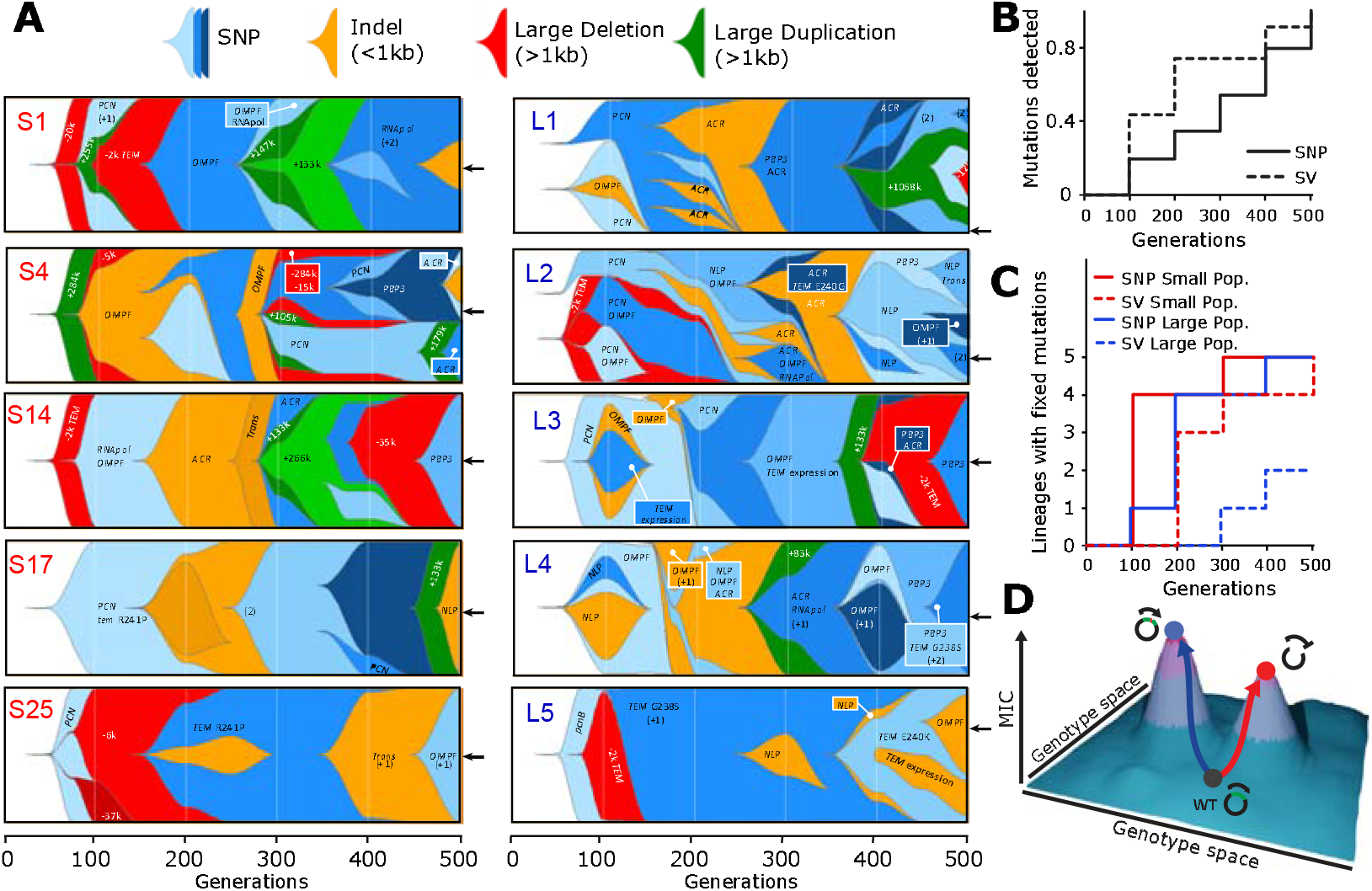
Temporal dynamics of genomic changes. (**A**) Muller plots inferred for five small and five large populations based on a comparison of population metagenomes at 100-generation intervals with final clone genotypes (indicated by arrows). Shown are mutations reaching at least 10% frequency and the functional targets they affect (see Fig. 3C), where applicable. (**B**) Time to first detection of SNPs and SVs for the 10 populations combined. (**C**) Time to fixation of SNPs and SVs for small and large populations separately. (**D**) Cartoon showing the adaptive consequences of two common trajectories (Fig. S19), either involving the high-rate deletion of TEM, which is twice as common in small populations, leading to modest CTX resistance levels (red), or large-benefit activation of TEM common in large populations and leading to significantly higher CTX resistance levels (blue) (Fig. S20).

Second, we used Wright-Fisher simulations to estimate the mutation rates and fitness effects of SNPs, indels and SVs that best explain their observed frequencies in the 91 non-mutator clones, assuming exponentially distributed and non-epistatic mutation effects [13]. This yielded selection coefficients of 0.41 for SNPs, 0.25 for indels and IS-element insertions and 0.14 for SVs, with corresponding mutation rates of 2.2 × 10^−8^ for SNPs, 1.8 × 10^−7^ for indels and IS-element insertions and 7.1 × 10^−6^ for SVs, in support of our hypothesis. The relative selection coefficients of the three mutation classes were confirmed by estimating their effects on the observed MIC values of these clones using a general linear model, which were approximately 2.5-fold larger for SNPs than SVs (Table S12).

Finally, we had a closer look at the common SV and SNP affecting β-lactamase TEM-1. Using pairwise competition assays [13], we found that the deletion of *bla*_TEM_ which was twice as common in small than in large populations was nearly neutral (Fig. S3), indicating that it was driven by its high mutation rate alone. In contrast, activating SNP G238S, expected to occur at a much lower rate, had a substantial benefit under the selective conditions (Fig. S3), explaining its role in rescuing TEM in 54% of the large and none of the small populations (for example, see population L5 in Fig. 4A). To examine the adaptive consequences of these mutations, we analyzed trajectories based on associations between mutations in different functional targets [13]. Small populations showed no clear associations, whereas large populations used two alternative trajectories: one combining the deletion of TEM with target alteration and efflux upregulation, and another involving TEM activation, downregulation of plasmid copy number and TEM expression and upregulation of outer-membrane vesicles (Fig. S19). Importantly, populations activating TEM reach higher resistance levels than those deleting TEM, particularly large populations (*P*<0.0001, Fig. S20), indicating that these trajectories have distinct adaptive consequences (Fig. 4D).

The observed tradeoff between rate and fitness effects among two major classes of mutations strengthens the impact of the population size on the contribution of high-rate and large-benefit mutations. Clearly, the adaptive consequences of high-rate versus large-benefit mutations also depend on the type of mutations. While in our study high-rate deletions of the β-lactamase constrained adaptation by preventing SNPs to activate the enzyme against the antibiotic, high-rate gene amplifications may, in contrast, facilitate adaptation through enhanced survival under stress [23] or increased mutation supplies and evolvability [22]. The interaction between high-rate SVs and large-benefit SNP is highly relevant for the clinical evolution of antibiotic resistance, since antibiotic-resistance genes are often flanked by repeat sequences facilitating their rapid deletion or amplification [24].

## Acknowledgments

We thank Bertha Koopmanschap (deceased) for practical help, and Duur Aanen, Su-Chan Park and Michael Lässig for discussion.

## Author contribution

Conceptualization: MFS and JAGMdeV; experimental design: MFS, JK and JAGMdeV; experiments: MFS, MPZ and PR; analysis: MFS, MPZ, SH, PR, ES, JK and JAGMdeV; writing: MFS, MPZ, PR, JK and JAGMdeV.

## Data and materials availability

Data described in the paper are presented in the supplementary materials. Raw data will be made available on Dryad upon publication.

## Literature

1. de Visser, J.A.G.M and J. Krug, Empirical fitness landscapes and the predictability of evolution. Nature Reviews Genetics, 2014. 15(7): p. 480–490.

2. Lässig, M., V. Mustonen, and A.M. Walczak, Predicting evolution. Nature Ecology & Evolution, 2017. 1: p. 0077.

3. Mas, A., Y. Lagadeuc, and P. Vandenkoornhuyse, Reflections on the Predictability of Evolution: Toward a Conceptual Framework. iScience, 2020. 23(11): p. 101736.

4. Palmer, A.C. and R. Kishony, Understanding, predicting and manipulating the genotypic evolution of antibiotic resistance (Progress). Nature Reviews Genetics, 2013. 14: p. 243–248.

5. Sommer, M.O.A., et al., Prediction of antibiotic resistance: time for a new preclinical paradigm? Nature Reviews Microbiology, 2017. 15: p. 689.

6. Baym, M., et al., Spatiotemporal microbial evolution on antibiotic landscapes. Science, 2016. 353(6304): p. 1147–1151.

7. Good, B.H., et al., The dynamics of molecular evolution over 60,000 generations. Nature, 2017. 551: p. 45.

8. Tenaillon, O., et al., The Molecular Diversity of Adaptive Convergence. Science, 2012. 335: p. 457–461.

9. Bailey, S.F., et al., What drives parallel evolution? BioEssays, 2017. 39(1): p. e201600176.

10. Garoff, L., et al., Population Bottlenecks Strongly Influence the Evolutionary Trajectory to Fluoroquinolone Resistance in Escherichia coli. Molecular Biology and Evolution, 2020. 37(6): p. 1637–1646.

11. Storz, J.F., Causes of molecular convergence and parallelism in protein evolution. Nature Reviews Genetics, 2016. 17(4): p. 239–250.

12. Orr, H.A., The probability of parallel evolution. Evolution, 2005. 59: p. 216–220.

13. Materials and methods are available as suporting material on Science online.

14. Gerrish, P.J. and R.E. Lenski, The fate of competing beneficial mutations in an asexual population. Genetica, 1998. 102/103: p. 127–144.

15. Good, B.H., et al., Distribution of fixed beneficial mutations and the rate of adaptation in asexual populations. Proceedings of the National Academy of Sciences USA, 2012. 109: p. 4950–4955.

16. Gomez, K., J. Bertram, and J. Masel, Mutation bias can shape adaptation in large asexual populations experiencing clonal interference. Proceedings of the Royal Society B: Biological Sciences, 2020. 287(1937): p. 20201503.

17. Payne, J.L., et al., Transition bias influences the evolution of antibiotic resistance in Mycobacterium tuberculosis. PLOS Biology, 2019. 17(5): p. e3000265.

18. Sackman, A.M., et al., Mutation-Driven Parallel Evolution during Viral Adaptation. Molecular Biology and Evolution, 2017. 34(12): p. 3243–3253.

19. Stoltzfus, A. and D.M. McCandlish, Mutational Biases Influence Parallel Adaptation. Molecular Biology and Evolution, 2017. 34(9): p. 2163–2172.

20. Svensson, E.I. and D. Berger, The Role of Mutation Bias in Adaptive Evolution. Trends in Ecology & Evolution, 2019. 34(5): p. 422–434.

21. Salverda, M.L.M., J.A.G.M. de Visser, and M. Barlow, Natural evolution of TEM-1 beta-lactamase: experimental reconstruction and clinical relevance FEMS Microbiology Reviews, 2010. 34: p. 1015–1036.

22. Andersson, D.I. and D. Hughes, Gene Amplification and Adaptive Evolution in Bacteria. Annual Review of Genetics, 2009. 43: p. 167–195.

23. Nicoloff, H., et al., The high prevalence of antibiotic heteroresistance in pathogenic bacteria is mainly caused by gene amplification. Nature Microbiology, 2019. 4(3): p. 504–514.

24. Sandegren, L. and D.I. Andersson, Bacterial gene amplification: implications for the evolution of antibiotic resistance. Nature Reviews Genetics, 2009. 7: p. 578–588.

