## Supplemental Materials for "Population size mediates the contribution of high-rate and large-benefit mutations to parallel evolution"

### Table of contents


Table of Statistical Results 4

1. Clonal interference, mutation bias and population size 6

2. Experimental methods and results 9
*Media and bacterial strains* 9
*Evolution experiment* 9
*Resistance assays* 11
*Competition assays* 11
*TEM $\beta$-lactamase activity assays* 13
*Genomic DNA extraction* 14
*High throughput sequencing* 15

3. Genomic analyses 16
*Mutation classes* 16
*Primary analysis with a custom pipeline* 16
*Primary analysis with CLC Genomics Workbench* 16
*Combining results from the two analyses* 17
*Detection of structural variants (SVs)* 17
*Detection of mutations for population samples* 18

4. Identification of mutator clones 19

5. Overview of genomics data 20

6. dN/dS and dI/dS analysis 25

7. Repeatability of genomic changes 26
*Methodology* 26
*Nucleotide-level repeatability results* 26
*Gene-level repeatability results* 27

8. Regression analysis of the relative frequency of SVs 31

9. Dynamics of genomic changes 32
*Muller plots* 32
*Repeatability of evolution* 32

10. Inference of mutation parameters from Wright-Fisher simulations 35

11. MIC effects of different mutation classes 38

12. Analysis of functional targets and evolutionary trajectories 41
*Analysis of evolutionary trajectories* 45

### Index of Supplementary Tables and Figures


Table S1: Table of statistical results 4

Table S2: Overview of conditions for experimental treatments 11

Table S3: Fitness effects of common TEM deletion and activating mutation G238S 12

Table S4: Common deletions in evolved clones 24

Table S5: Common duplications in evolved clones 24

Table S6: dN/dS and dI/dS results 25

Table S7: Nucleotide-level *H*-indexes 28

Table S8: Gene-level *H*-indexes 30

Table S9: Comparison of mutations detected in clones and final populations 33

Table S10: Inferred mutation rates and selection coefficients 37

Table S11: Model selection with the AIC for mutational effect size models 39

Table S12: Estimated model parameters for mutational effect size models 40

Table S13: Overview of all mutations affecting multiple-hit genes 44

Fig. S1: Condition at which mutation and selection bias are balanced 8

Fig. S2: CTX concentration over generations in the evolution experiment 10

Fig. S3: Relative fitness of common mutations affecting TEM1 13

Fig. S4: TEM β-lactamase activity measurements 14

Fig. S5: Histogram of the frequency of mutations in the chromosome 17

Fig. S6: Histogram of the number of mutations per clone for all 112 populations 19

Fig. S7: Mutation frequencies per mutation (SNP, Indel, SV) class for non-mutator clones 20

Fig. S8: Total mutation frequencies per clone per treatment 21

Fig. S9: Mutation frequencies per mutation class and per treatment 21

Fig. S10: Mutation frequencies per mutation class per chromosome 22

Fig. S11: Comprehensive overview of mutations for all 112 populations 23

Fig. S12: Nucleotide-level pairwise similarity index (*H*-index) 27

Fig. S13: Comparison of the nucleotide-level pairwise similarity indexes 29

Fig. S14: Gene-level pairwise similarity index (*H*-index) 29

Fig. S15: Regression analysis of fraction of SVs 31

Fig. S16: Similarity indexes (*H*-indexes) for time-course meta-population samples 34

Fig. S17: Comparison optimized Wright-Fischer model in comparison to the experimental data 37

Fig. S18: Overview of the eight different mutational effect-size models fitted to the MIC data 39

Fig. S19: Associations between functional targets with frequent mutations 45

Fig. S20: MIC doublings for genotypes that activate, delete or maintain TEM1 46

### Table of Statistical Results

**Table S1:** Statistical results from tests of data presented in the figures.

| Figure | Comparison | Test | Results |
|---|---|---|---|
| 2A | Minimal inhibitory concentration (MIC) of all large vs. small populations | MWU[a] | $U = 56.5$, $N = 96$, $P < 0.001$ |
| 2B | Total number of mutations in CTX non-mutator populations (large and small) vs. control populations with no CTX | MWU[a] | $U = 1352.5$, $N = 107$, $P < 0.0001$ |
| 2B | Total number of mutations in large vs. small non-mutator populations | MWU[a] | $U = 823$, $N = 107$, $P = 0.405$ |
| 2B | SNPs in large vs. small non-mutator populations | MWU[a] | $U = 1186.5$, $N = 91$, $P < 0.0001$ |
| 2B | SNPs in large vs. small non-mutator populations; plasmid only | MWU[a] | $U = 1196$, $N = 91$, $P < 0.0001$ |
| 2B | SNPs in large vs. small non-mutator populations; chromosome only | MWU[a] | $U = 920.5$, $N = 91$, $P = 0.076$ |
| 2B | SVs in large vs. small non-mutator populations | MWU[a] | $U = 252$, $N = 91$, $P < 0.0001$ |
| 2B | SVs in large vs. small non-mutator populations; plasmid only | MWU[a] | $U = 511$, $N = 91$, $P = 0.0149$ |
| 2B | SVs in large vs. small non-mutator populations; chromosome only | MWU[a] | $U = 288.5$, $N = 91$, $P < 0.0001$ |
| 2B | Indels in large vs. small non-mutator populations | MWU[a] | $U = 707.5$, $N = 91$, $P = 0.794$ |
| 3A | Nucleotide-level pairwise repeatability vs. gene-level repeatability, for all non-mutator large and small populations | MWU[a] | $U = 10{,}828$, $N = 107$, $P < 0.0001$ |
| 3A | Gene-level pairwise repeatability in small vs. large non-mutator populations | MWU[a] | $U = 149$, $N = 91$, $P < 0.0001$ |
| 3A | Nucleotide-level pairwise repeatability in small vs. large non-mutator populations | MWU[a] | $U = 507$, $N = 91$, $P = 0.032$ |
| 3B | Nucleotide-level pairwise repeatability in small vs. large non-mutator populations, SNPs only | MWU[a] | $U = 13$, $N = 91$, $P < 0.0001$ |
| 3B | Nucleotide-level pairwise repeatability in small vs. large non-mutator populations, SVs only | MWU[a] | $U = 1335$, $N = 91$, $P < 0.0001$ |
| 3B | Nucleotide-level pairwise repeatability in small vs. large non-mutator populations, Indels only | MWU[a] | $U = 979$, $N = 91$, $P = 0.017$ |

[a] Mann-Whitney *U* test.

**Table S1 (Continued):** Statistical results from tests presented in the figures.

| Figure | Comparison | Test | Results |
|---|---|---|---|
| 3B | Nucleotide-level pairwise repeatability in small vs. large non-mutator populations, SNPs in plasmid only | MWU[a] | $U$ = 300, $N$ = 91, $P$ < 0.0001 |
| 3B | Nucleotide-level pairwise repeatability in small vs. large non-mutator populations, SNPs in chromosome only | MWU[a] | $U$ = 271, $N$ = 91, $P$ < 0.0001 |
| 3B | Nucleotide-level pairwise repeatability in small vs. large non-mutator populations, SVs in plasmid only | MWU[a] | $U$ = 1054, $N$ = 91, $P$ < 0.001 |
| 3B | Nucleotide-level pairwise repeatability in small vs. large non-mutator populations, SVs in chromosome only | MWU[a] | $U$ = 1308, $N$ = 91, $P$ < 0.0001 |
| 4B | Time until detection of SVs vs. SNPs | $\chi^2$ test | $\chi^2$ = 4.13, $P$ = 0.042 |
| 4C | Time until fixation of SVs in small vs. large populations | $\chi^2$ test | $\chi^2$ = 5.975, $P$ = 0.015 |
| S14B | Gene-level pairwise repeatability in small vs. large populations, SNPs only | MWU[a] | $U$ = 163, $N$ = 91, $P$ < 0.0001 |
| S14B | Gene-level pairwise repeatability in small vs. large populations, SVs only | MWU[a] | $U$ = 1336, $N$ = 91, $P$ < 0.0001 |
| S14B | Gene-level pairwise repeatability in small vs. large populations, Indels only | MWU[a] | $U$ = 602.5, $N$ = 91, $P$ = 0.199 |
| S20 | CTX MIC of clones deleting and activating TEM (small and large populations combined) | MWU[a] | $U$ = 762.5, $N$ = 56, $P$ < 0.0001 |

[a] Mann-Whitney *U* test.

### 1. Clonal interference, mutation bias and population size

In large asexual populations, simultaneously segregating mutations compete for fixation, a phenomenon generally referred to as clonal interference [1-3]. The competition between beneficial mutations implies that mutations of strong effect fix preferentially, but interference also modifies the fixation of linked mutations that are neutral or weakly deleterious [4, 5].

As a simple scenario that illustrates how the balance between mutation supply and selection changes with population size, we consider two competing beneficial mutations with rates $\mu_1 > \mu_2$ and selection coefficients $s_1 < s_2$ that arise in a monomorphic population. For small populations in the strong selection/weak mutation (SSWM) regime, mutations of type $i = 1{,}2$ appear at rate $N\mu_i$ and fix with a probability proportional to $s_i$, where $N$ denotes the population size and the SSWM conditions imply that $N\mu_i \ll 1$ and $Ns_i \gg 1$ [6]. The rate of fixation is therefore proportional to $s_i\mu_i N$, and a straightforward calculation shows that the probability that the mutation 2 of stronger effect fixes first is [7]:

$$P_2 = \frac{\mu_2 s_2}{\mu_1 s_1 + \mu_2 s_2}. \tag{3}$$

The condition for selection and mutation bias to balance thus reads $s_1\mu_1 = s_2\mu_2$ in the SSWM regime.

An approximate expression for $P_2$ that remains valid also in the presence of a moderate amount of clonal interference can be obtained by adapting the approach of [8]. The original calculation considered the special case when $\mu_1 = 2\mu_2$ and was based on an estimate of the probability that the mutation of weaker effect fixes before the strong-effect mutation has been established. Generalizing the argument to arbitrary mutation rates and including a heuristic modification that enforces the constraint that $P_2 = \frac{\mu_2}{\mu_1+\mu_2}$ when $s_1 = s_2$, one arrives at the expression:

$$P_2 = 1 - \frac{\mu_2 s_2}{\mu_1 s_1 + \mu_2 s_2} exp\left\{-2N\ln(Ns_1)\mu_2\left[\frac{s_2}{s_1} - 1\right]\right\}. \tag{4}$$

This reduces to (3) for small population sizes. Simulations presented in [8] show that this approach is rather accurate at least for the special case $\mu_1 = 2\mu_2$, but more recent work has revealed that it breaks down when the mutation supply rates $N\mu_1$, $N\mu_2$ become large. A formula similar to (4) was presented by Svensson and Berger [9]. **Fig. 1** in the main text illustrates the dependence of $P_2$ on the selection bias $s_2/s_1$ for fixed mutation bias $\mu_1/\mu_2 = 100$ and different population sizes.

To address the regime of strong clonal interference, we refer to a recent study of Gomez et al. [10] on the competition between two simultaneously adapting traits, characterized by different selection coefficients and mutation rates. The traits correspond to the different mutation classes in our system, and the main conclusion that is relevant in our context is that adaptation is dominated by the trait (mutation class) that has the faster rate of adaptation in isolation. Theory reviewed in [3] shows that, to leading order for very large $N$, the rate of adaptation $v$ for a population subject to a single class of beneficial mutations with selection coefficient $s$ occurring at rate $U$ is given by:

$$v = \frac{s^2 \ln N}{\ln^2 U}. \quad (5)$$

Note that here $U \ll 1$ denotes the *total* rate of beneficial mutations per individual and generation. In contrast to the SSWM regime where the rate of adaptation is proportional to the mutation supply rate $NU$, under conditions of strong clonal interference the dependence on $U$ is only logarithmic. As a consequence, moderate differences in selection strength between different mutation classes can be overcome by mutation rate difference only if the mutation bias is extremely strong. Specifically, considering two classes characterized by parameters $U_1 > U_2$ and $s_1 < s_2$, according to Eq. (5), the corresponding rates of adaptation are equal when

$$U_1 = (U_2)^{s_1/s_2}. \quad (6)$$

For example, to overcome a selection bias $s_2/s_1 = 2$ the mutation rate has to be elevated from $U_2$ to $U_1 = \sqrt{U_2} \gg U_1$. The relation (6) is illustrated in **Fig. S1**.

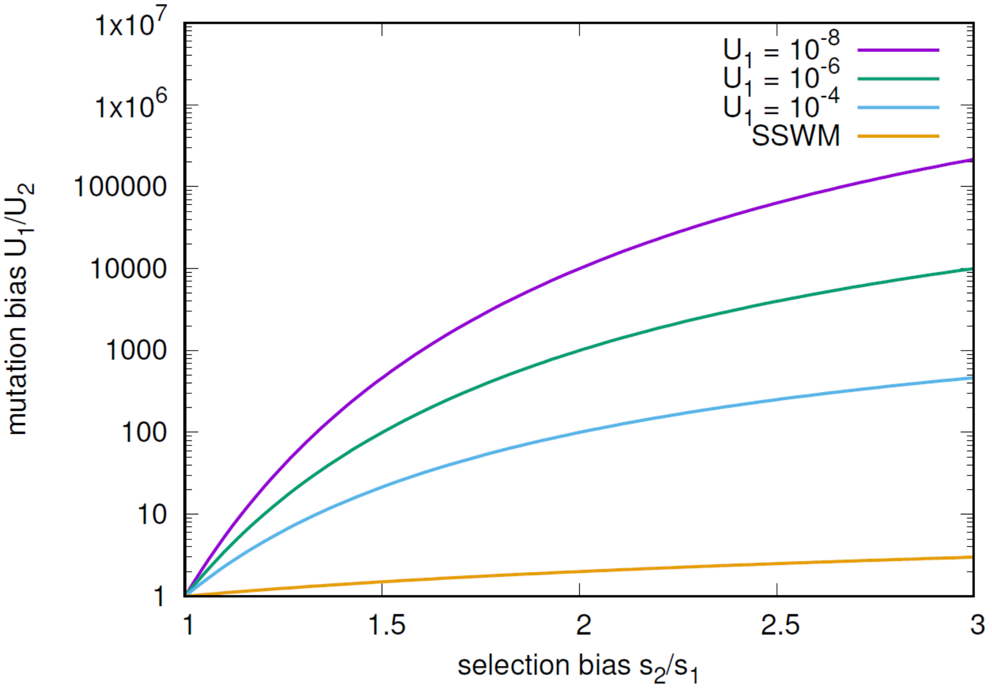


**Fig. S1**. Illustration of condition (4) at which mutation and selection bias are balanced in the regime of very strong clonal interference.

#### 2. Experimental methods and results

##### *Media and bacterial strains*

For all experiments, we used a modified Luria broth (LB), which here is 10 g/L trypticase peptone, 5 g/L yeast extract and 5 g/L NaCl. For plates, 15 g/L agar was added. *Escherichia coli* strains REL606 (Ara-) and REL607 (Ara+) [11] were used for all experiments.

##### *Evolution experiment*

We electro-transformed REL606 and REL607 cells derived from a single colony with the pACTEM1 plasmid expressing TEM1 β-lactamase [12] and subsequently plated them on LB supplemented with 15 µg/mL tetracycline for selection of transformants. A different colony was used to initiate 72 small, 24 large and 16 control population (large populations with no antibiotics or tetracycline only, N = 8 each). Colonies were grown up o/n in 1 mL LB with 15 µg/mL tetracycline. These overnight cultures were then used to initiate the serial passaging experiment with a 1:1,000 dilution in LB with different supplements depending on the treatment (**Table S2**), including 0.011 µg/mL cefotaxime (CTX), 50 µM isopropyl β-D-1-thiogalactopyranoside (IPTG) to induce TEM expression, and 15 µg/mL tetracycline to force plasmid maintenance (except for control populations C1-8). Small populations were represented by 200 µl cultures in 300 µl flat-bottom wells, distributed in a checkerboard pattern across two 96-well microtiter plates (ThermoFisher Nunc): Ara+ and Ara- populations were alternated and wells in between bacterial cultures were filled with sterile medium to control for potential contamination. Large populations were represented by 20 mL cultures in 50 mL tubes (Greiner), and were also transferred in alternating fashion with respect to Ara-marker. All cultures were kept at 37 °C with agitation (220 rpm).

Transfers involved daily 1:1,000 dilutions (volume:volume) for 50 subsequent days (equivalent to ~500 bacterial generations). In the absence of CTX, these conditions yielded effective population sizes of ~2 x $10^6$ and ~2 x $10^8$ for small and large populations, respectively. These population sizes were chosen based on the expectation of substantial differences in the strength of clonal interference from previous work [13]. To maximize selection for CTX resistance, we increased the concentration of CTX as follows. Before transfer, the optical density ($OD_{600}$) was measured with a Victor$^3$ plate reader (Perkin Elmer). When the $OD_{600}$ was at least 75% of that of the ancestral strain in the absence of CTX, the concentration of CTX was increased by a factor of $2^{0.25}$ (~19%), so that after four increases the concentration doubled (**Fig. S2**). In rare cases when the $OD_{600}$ dropped below 25% of its maximum, the concentration of CTX was decreased by a factor of $2^{0.25}$. After 10, 20, 30, 40 and 50 transfers, samples were plated on TA agar to check for Ara-marker, and glycerol stocks of the populations were prepared and stored at -80 ˚C.

On three occasions, growth in control wells with medium was detected. In those cases, the entire plate was restarted from the previous day's plate, which was stored at 4˚C. On two occasions the wrong

Ara-marker type was detected in a small population, and those populations were restarted from frozen samples from the previous time point. The final cultures were plated out on LB agar with 15 μg/mL tetracycline, and a random clone was selected after overnight growth for subsequent analysis and grown up in 1 mL LB with 15 μg/mL tetracycline to prepare glycerol stocks, except in the case of populations C1-C8, for which no antibiotics were used.

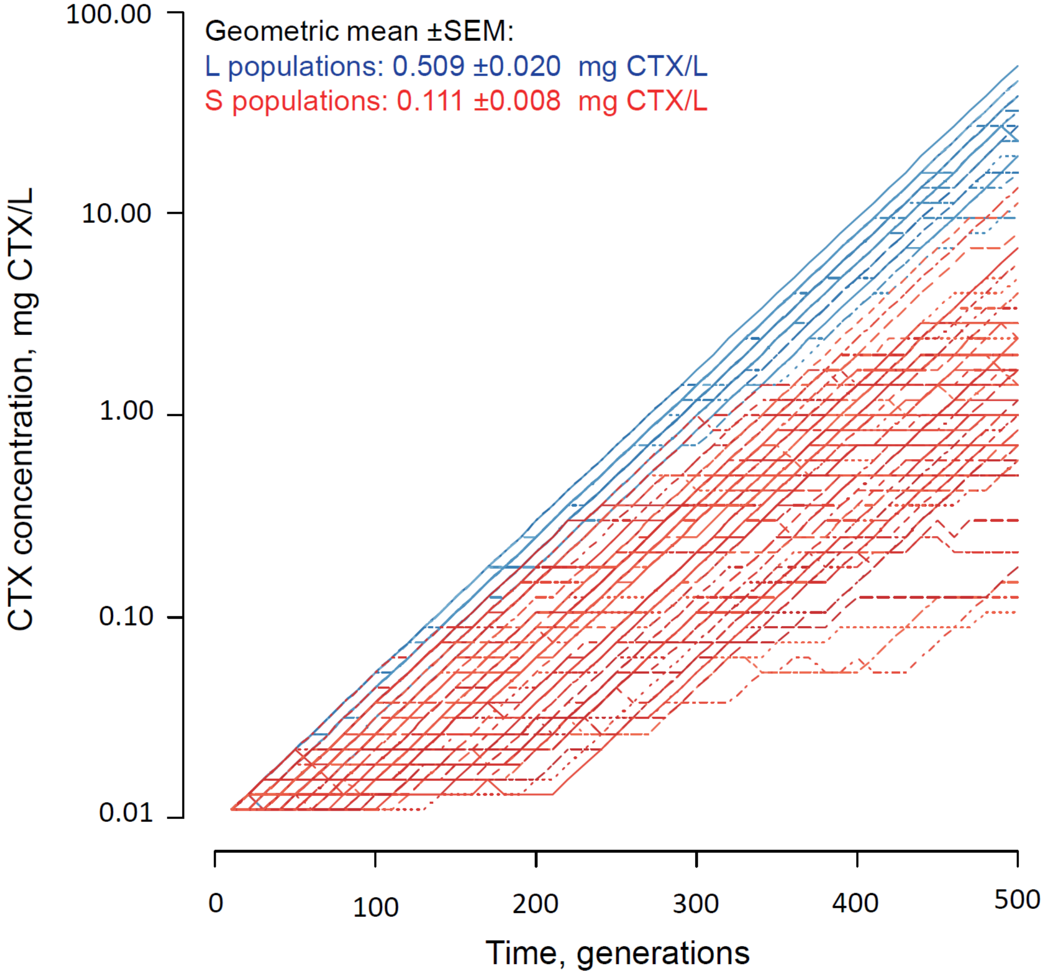


**Fig. S2:** CTX concentrations during the evolution experiment based on daily $2^{0.25}$-fold increases when the $OD_{600}$ was higher than 75% of the ancestral value without CTX (see text). Red lines represent small populations (S), blue lines large populations (L). Shades and line types have been varied randomly to better distinguish replicate populations.

**Table S2**. Overview of conditions for experimental treatments.

| Condition | Pop. Numbers[a] | Replicates | Culture volume (mL) | CTX (µg/mL) | IPTG (µM) | Tetracycline (µg/mL) |
|---|---|---|---|---|---|---|
| Small | S1-S72 | 72 | 0.2 | ≥ 0.011 | 50 | 15 |
| Large | L1-L24 | 24 | 20 | ≥ 0.011 | 50 | 15 |
| No antibiotic Control | C1-C4 | 4 | 20 | 0 | 50 | 0 |
| | C5-C8 | 4 | 20 | 0 | 0 | 0 |
| Tetracycline only Control | C9-C12 | 4 | 20 | 0 | 50 | 15 |
| | C13-C16 | 4 | 20 | 0 | 0 | 15 |

[a] Note that in some of the primary analysis files and scripts the populations are numbered consecutive from 1-112. In these cases, 1-24 = L1-L24, 25-96 = S1-S72, 97-112= C1-C16.

#### *Resistance assays*

Minimal inhibitory concentration (MIC) assays were performed to determine the resistance of clones and population samples from the final time point, using an assay representative of the experimental conditions. Cultures with a total volume of 200 µL LB were setup, including with $5 \times 10^5$ cells/mL, 50 µM IPTG, 15 µg/mL tetracycline, and two-fold serial dilutions of CTX ranging from 512 to 0.25 µg CTX/mL, as well as no CTX. These cultures were grown for 24 h at 37 °C with agitation. Growth was then determined by observing an $OD_{600} > 0.075$, whilst all wells for which $0.04 < OD_{600} < 0.075$ were inspected visually to determine growth. Three biological replicates were performed.

#### *Competition assays*

We measured the fitness consequences of two common mutations affecting TEM-1 β-lactamase, i.e. the deletion of ~2 kbp from pACTEM, including *bla*$_{\text{TEM-1}}$ and its repressor *lacI*, and activating SNP G238S, by running pairwise competition assays with constructed mutants against ancestral strain REL606/pACTEM1. To do so, we transformed YFP- and CFP-labelled versions of REL606 with ancestral pACTEM1, pacDEL and pACTEM-G238S plasmids, which were isolated from evolved clones. The YFP- and CFP-labelled REL606 strains were provided by Vaughn Cooper (University of Pittsburgh). Competition assays were performed in 96-well microtiter plates containing 200µl LB supplemented with 50 µM IPTG, 15 µg/mL tetracycline and 0, 0.01 and 0.04 mg CTX/L and were initiated with 1:1,000 dilution from overnight cultures. Each competition was performed with six-fold replication in both fluorescently-marked backgrounds (12-fold replication in total). The frequency of both competitors was estimated prior to inoculation (t=0h) and after incubation (t=24h) at 37°C with shaking (600rpm) by counting YFP and CFP cells in samples of ~10,000 cells using a flow cytometer (MacsQuant Analyzer 10, Miltenyi). Prior to flow cytometry, cells were washed twice and resuspended in 200µl phosphate-buffered saline. Relative fitness was estimated by the ratio of Malthusian parameters [11], and tested for significant deviation from 1 using one-sample *t*-tests with serial-Bonferroni correction for multiple testing (**Table S3**, **Fig. S3**). The estimates indicate that the common deletion of TEM and repressor LacI from the plasmid has no fitness consequences, while common activating mutation G238S provides a benefit at CTX concentrations above the starting concentration of 0.011 mg CTX/L (but has a cost below that).

**Table S3**. Relative fitness estimates of common TEM deletion and TEM-activating SNP G238S from pairwise competitions against ancestral strain REL606/pACTEM1 for three CTX concentrations.

| | TEM deletion | | | G238S | | |
|---|---|---|---|---|---|---|
| mg CTX/L | 0 | 0.01 | 0.04 | 0 | 0.01 | 0.04 |
| | 0.8794 | 0.8272 | 1.3131 | 0.8022 | 1.1772 | 1.1961 |
| | 0.9328 | 1.0693 | 1.0331 | 0.8740 | 1.3671 | 1.7191 |
| | 1.1570 | 0.9719 | 1.0866 | 0.6502 | 0.9620 | 1.2720 |
| | 0.8059 | 0.8770 | 0.8948 | 0.7498 | 0.9814 | 1.6123 |
| | 0.8668 | 0.8940 | 0.8248 | 0.6856 | 1.0569 | 1.2841 |
| | 0.9560 | 0.9389 | 0.8558 | 0.6772 | 1.0220 | 1.1632 |
| | 1.1081 | 1.6587 | 1.7013 | 0.8517 | 0.8926 | 1.1109 |
| | 1.2035 | 1.0269 | 0.9855 | 0.8426 | 0.8736 | 0.8826 |
| | 1.2748 | 0.8306 | 0.9324 | 0.5443 | 0.8073 | 1.1904 |
| | 0.9590 | 1.2066 | 0.7801 | 0.8856 | 1.0843 | 1.5536 |
| | 0.9732 | 0.6188 | 0.7894 | 0.7701 | 0.8711 | 1.1769 |
| | 1.1617 | 0.5935 | 0.6507 | 0.6619 | 1.1114 | 1.0462 |
| Mean | 1.0232 | 0.9595 | 0.9873 | 0.7496 | 1.0173 | 1.2673 |
| S.E.M. | 0.0437 | 0.0808 | 0.0816 | 0.0308 | 0.0451 | 0.0705 |
| *t* (versus 1) | 0.5306 | -0.5017 | -0.1558 | -8.1273 | 0.3823 | 3.7896 |
| Two-tailed *P* | 0.6062 | 0.6258 | 0.8790 | 5.62E-06* | 0.7095 | 0.0030* |

*: significant after serial-Bonferroni correction for multiple testing

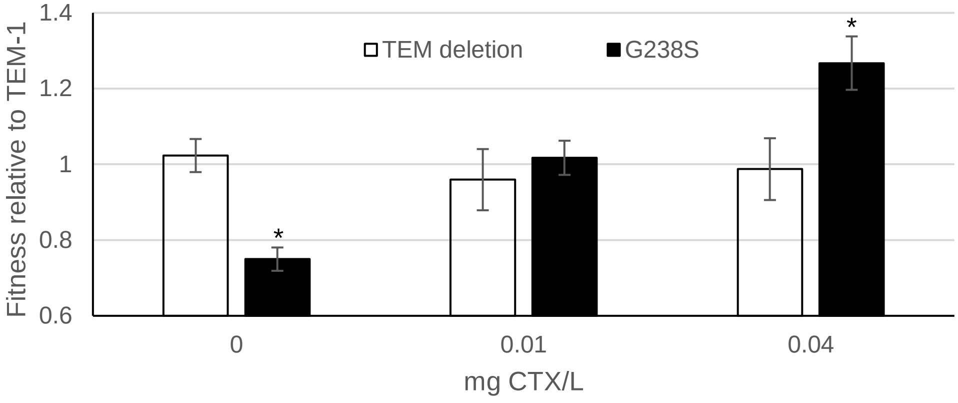


**Fig. S3:** Relative fitness effect of common ~2kbp TEM/lacI deletion and most common TEM-activating SNP (G238S), measured in pairwise competitions against ancestor TEM1 using fluorescently-labelled strains in the presence of varying concentrations of CTX. An asterisk indicates a significant difference between the TEM deletion or G238S versus TEM1 (see **Table S3)**.

#### *TEM β-lactamase activity*

To understand the effects of common chromosomal mutations in genes *pcnB* and *polA* on plasmid copy number, their effect on TEM expression was measured in the large populations. First, each final clone of the large populations was cured of the pACTEM plasmid. This was done by performing two passages with 50 µM IPTG but without any antibiotics, followed by plating of the populations and screening for clones without tetracycline resistance. The cured clones were then electro-transformed with the ancestral pACTEM1 plasmid. For each final clone, a transformant carrying pACTEM1 was then grown up overnight in LB with 15 µg/mL tetracycline, and subsequently diluted 1,000-fold into LB with 15 µg/mL tetracycline and 50 µM IPTG. After 2 h incubation, the culture was diluted 10-fold into ice-cold phosphate buffered saline (PBS) pH = 7.4 with 2 mg/mL nitrocefin. By pelleting cells (5 min. at 5000 rcf at 4 °C), we could assay the supernatant or the resuspended cells for TEM activity [11]. The plate was then incubated at room temperature (21 °C) for 1 h, and the $OD_{490}$ was measured every minute using a Victor$^3$ plate reader (Perkin-Elmer). We included a standard curve, based on a stationary phase culture diluted by 4-fold steps in LB. For each time point, we fitted a linear regression to the square root of the expected relative activity (as determined by the dilution of the stationary phase sample) and the observed activity. We then chose the time point with the highest coefficient of determination ($r^2$) for the standard curve and used only the data from this time point for determining relative TEM activity in the experimental samples using the fitted linear relationship. We then normalized TEM activity by the $OD_{600}$ value of the culture prior to dilution into PBS, to take into account minor differences in cellular density. Three biological replicates were analyzed for each final clone.

As each evolved final clone from the large populations had the ancestral TEM plasmid reintroduced into it, we measured how mutations in *pcnB* and *polA* affected TEM beta-lactamase activity. We found that in most clones, beta-lactamase activity was reduced strongly (**Fig. S4**). This reduction of activity was statistically significant for all evolved clones except populations 5 and 18, as determined by a pairwise *t*-test with serial Bonferroni correction. Population 18 is the only of the 24 large populations that does not have mutations in the *pcnB* or *polA* genes, while population 5 harbors a 210-bp deletion in *pcnB*. Given the significant reduction of plasmid copy number in 22 of the 23 populations with mutations in *pcnB* or *polA* (of which many presumably inactivate the gene function), while the only population without these mutations shows no decrease, strongly suggested that mutations in *pcnB* and *polA* result in a reduction of plasmid copy number. As TEM expression is costly, these reduced expression and activity levels are likely to be beneficial for growth in the absence or at low concentrations of CTX.

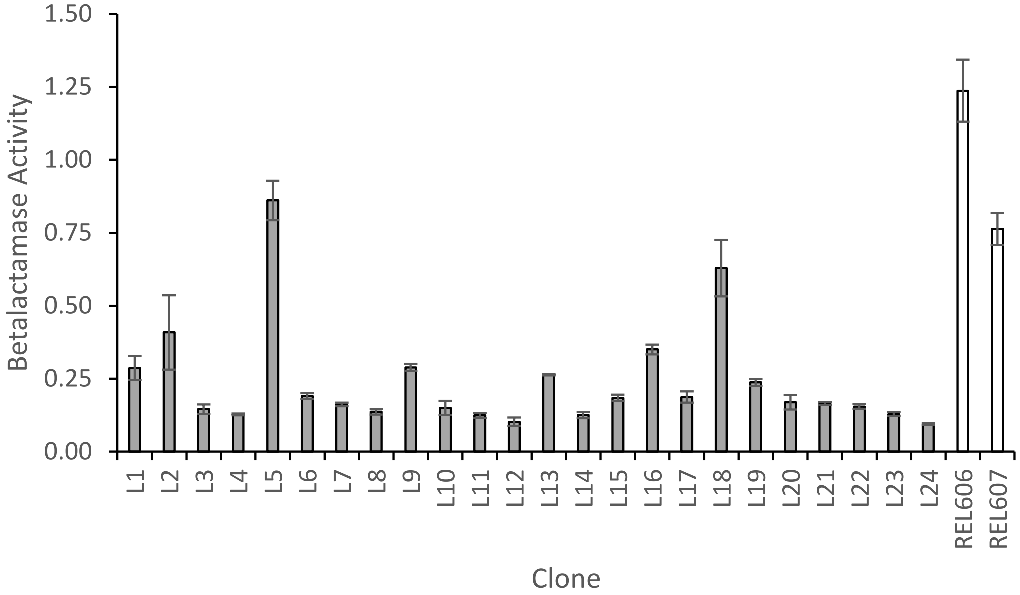


**Fig. S4:** TEM beta-lactamase activity measurements in supernatants from large populations with the ancestral pACTEM plasmid reintroduced. On the x-axis is the clone from the large populations as well as the two ancestors, and on the y-axis is the TEM activity, expressed relative to average of the ancestral REL606/607 clones. For most clones, β-lactamase activity was strongly reduced. All clones had significantly lower activity than both ancestral strains, except L5 and L18. There was a good agreement between measurements on supernatants and cells ($r^2$ = 0.988).

#### *Genomic DNA extraction*

We extracted genomic DNA from the randomly selected clone from the final time point of the small, large and control populations. In addition, from the pool of transformants from which clones were selected to found experimental populations, we sequenced two Ara+ and two Ara- clones. We also sequenced the metagenomes of five small and five large populations from five time points (100, 200, 300, 400 and 500 generations). The populations we sequenced were L1-L5, and S1, S4, S14, S17 and S25. For the small populations, we selected two populations with TEM-activating mutations in the final sequenced clone (S17 and S25) and three random populations. Total genomic DNA was isolated with the Gentra Puregene Yeast/Bacterial DNA extraction kit (Qiagen), following the manufacturer's instructions, including overnight incubation at room temperature with agitation for DNA resuspension. In addition, plasmid DNA was isolated separately with Nucleospin Plasmid DNA kit (Machery-Nagel) and then added to the genomic samples from the same clone/population, to ensure high coverage of the plasmid.

##### *High throughput sequencing*

The NexteraXT (Illumina) kit was used for library preparation, and libraries were sequenced by HiSeq 2500 PE150. Library preparation and sequencing were performed by the Cologne Center for Genomics (https://portal.ccg.uni-koeln.de/ccg/index.php). For the 112 clones from the final populations, coverage of the bacterial chromosome was 42.08 ±0.59 (mean ±SEM), and coverage of the plasmid was 524.70 ±66.96. Coverage calculations for the plasmid are based only on those clones with an intact plasmid, and not those with large deletions.

### 3. Genomic analyses

For detecting mutations in the sequenced clones, two different approaches were used for primary analysis of the Illumina data. First, the data were analyzed using a custom pipeline. Second, the data were analyzed with CLC Genomics Workbench v8.01 (Qiagen Bioinformatics). Data from the two approaches were then compared to validate the identification of genomic changes.

#### *Mutation classes*

Note that as in the manuscript proper, we refer to three classes of mutations here: (1) single-nucleotide polymorphisms (SNPs): the substitution on a single nucleotide. (2) Indels: insertion sequence (IS) insertion, and duplication, deletions or inversions with a length <1 Kbp, and (3) structural variants (SVs): duplications, deletions or inversions with a length >1 Kbp.

#### *Primary analysis with a custom pipeline*

Raw sequences reads were preprocessed by first removing potential sequencing adaptors using Cutadapt [14] and then trimming low quality regions with Trimmomatic [15]. The cleaned reads were mapped to the genomic sequence using Bowtie2 [16] with default parameters and only those alignments with mapping quality equal or larger than 20 were kept. The alignments were summarized using the mpileup function of the samtools program [17]. Custom python scripts were used to summarize the observed nucleotide frequencies at all genomic positions from the mpileup output.

#### *Primary analysis with CLC Genomics Workbench*

The Illumina data were also analyzed with CLC Genomics Workbench. The trim sequences tool was used to trim demultiplexed reads, with default settings except a Phred threshold of 30 (expected error ≤ 1/100). Broken pairs were not discarded because our coverage on the chromosome was not very high. The trimmed reads were mapped to the REL606 genome (Genbank NC_012967.1) and the pACTEM1 sequence (Genbank MN386081), using default settings and with random allocation of ambiguously mapping reads. The basic variant detector was then used to identify mutations, again using default settings. All mutations with a frequency above 0.01 (i.e., 1%) were called. For a small number of regions of the genome where it proved difficult to disentangle mutational events, we extracted the reads from small regions (< 1,000-bp long) and performed *de novo* assembly in CLC Genomics Workbench, using default settings.

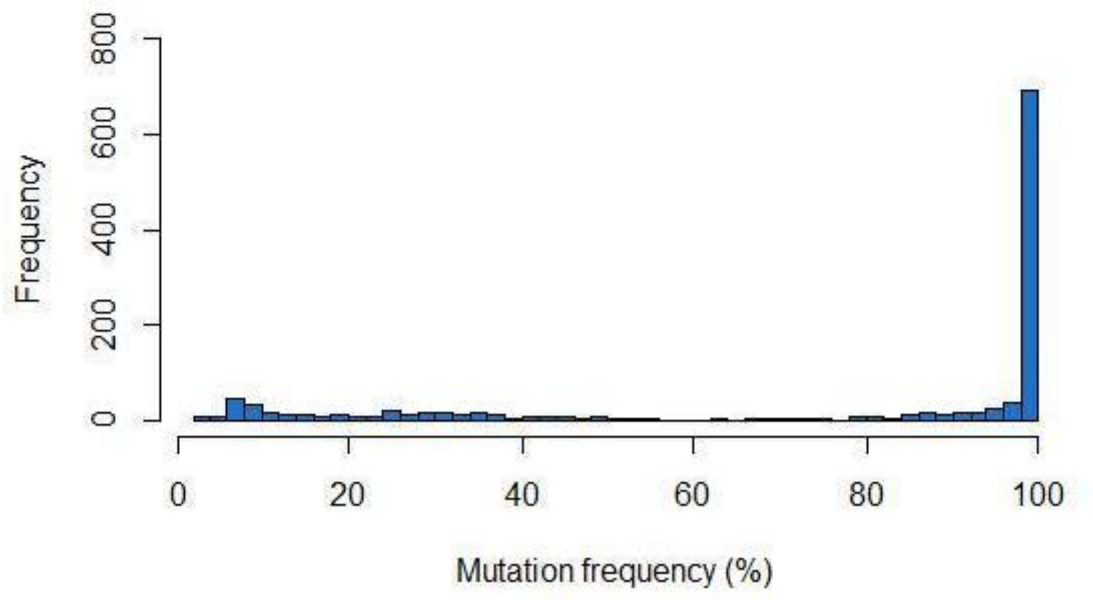


**Fig. S5:** Histogram of the frequency of mutations among reads mapped onto the chromosome for the CLC Genomics Workbench variant calling results of the sequenced clones, prior to manual curation.

##### *Combining results from the two analyses*

We compared the variants detected with these two approaches and combined them into a single database. We first eliminated all mutations (i) associated with the ancestral REL607 strain, and (ii) those that were detected in at least one of the four sequenced starting clones. We then flagged mutations that were (i) identified only by one sequence-analysis approach, or (ii) were not categorized as being fixed, or (iii) showed strong strand bias (< 40% or > 60% forward reads). We then manually curated all flagged mutations, discarding those mutations that appeared to be sequencing artifacts. There was generally good agreement between the two approaches, with 97.1% of the mutations accepted (after manual curation) being detected by both methods. The majority of mutations detected were present at high frequency in the reads, as even for the noncurated mutations 66.0 % were present at a frequency > 0.9 (**Fig. S5**).

##### *Detection of structural variants (SVs)*

During our manual curation of mutational events, we noticed that some larger Indels and SVs (SVs) – in particular events larger than the average read length – had not been detected. We therefore performed additional checks to detect these variants. First, we scanned the sequence alignments for low coverage regions (< 10 reads), and then manually mapped deletions spotted. Second, we plotted the coverage per base pair and in bins of 100 bp, over the entire genome. This allowed for quick but effective visual inspection, which we combined with *kmeans* algorithms to determine if loci with different levels of coverage were clustered. Upon visual inspection, any regions with higher coverage were then manually checked for duplications. For the analyses of clones, we expect duplications to have fold increases that are whole numbers. When increases in coverage were estimated – by dividing the coverage in the duplicated region by coverage in an immediately upstream region of length > 100,000 bp – only values above 1.75 were considered as real events. We only noticed two events with intermediate coverage levels (1.1 and 1.3 fold), which were disregarded. In addition, when manually curating events called by our custom pipeline and CLC Genomics Workbench, we noticed that some spurious events (e.g., low frequency SNPs) in fact indicated other mutational events, such as the addition of an IS element. Overall, all SVs (213) were detected through the procedures described here. For indels, about a quarter (26.1%, 70/268) were detected through the procedure described here or the general manual curation. Most SNPs were detected automatically, and only a small number was assigned after manual curation (3.6%, 26/706).

##### *Detection of mutations for population samples*

For detection of mutations in the population samples from different time points, we used CLC Genomics Workbench for primary sequence data analysis, as described above. Since we expected to find genetic variants at intermediate frequencies, we altered our criteria for accepting mutations. For SNPs and indels, ≥ 5 reads must be present for the mutation to be called, and it must reach a frequency ≥ 0.1 for at least one time point. Similarly, for the ancestral allele to be deemed present, it must also have ≥ 5 reads. Copy number variants must reach a frequency ≤ 0.75 or ≥ 1.25 at one time point to be deemed as real events. There are copies of *lacI* on both the bacterial chromosome and the pACTEM plasmid, which we could not distinguish due to ambiguity with read mapping.

### 4. Identification of mutator clones

To identify populations that evolved a mutator phenotype, we first looked for mutations in known mutator loci in the clones chosen for sequencing. We found non-synonymous mutations in *mutL* (population L15, S62) and *mutS* (populations L11, L21). We then considered the total number of mutations of all classes that had occurred per clone. Five populations were clearly outliers (**Fig. S6**), including the four populations with non-synonymous mutations in *mutL* (L15: 40 mutations; S65: 31 mutations) and *mutS* (L11: 68 mutations; L21: 42 mutations), but also including one other population without mutations in known mutator genes: S62 with 31 mutations. Given the high number of mutations that occurred in this clone, we considered this population also a putative mutator. Possibly its high number of mutations was due to unstable genetic changes, such as temporary copy number variation of sequences affecting DNA metabolism and repair. For all populations, the variation of the number of mutations per clone is much higher than its mean ($\mu$ = 9.611, $\sigma^2$ = 61.824) and the distribution is significantly different from a Poisson distribution (One-sample Kolmogorov-Smirnov test: $D$ = 1.890, $N$ = 112, $P$ = 0.002), whereas when we exclude these five putative mutators the variance is similar to the mean for the remaining populations ($\mu$ = 9.308, $\sigma^2$ = 10.574) and the distribution is not significantly different from a Poisson distribution ($D$ = 0.762, $N$ = 112, $P$ = 0.608). This result suggests the putative non-mutator populations are all acquiring mutations in a Poisson-like process at a similar rate, reinforcing our selection of mutator and non-mutator clones.

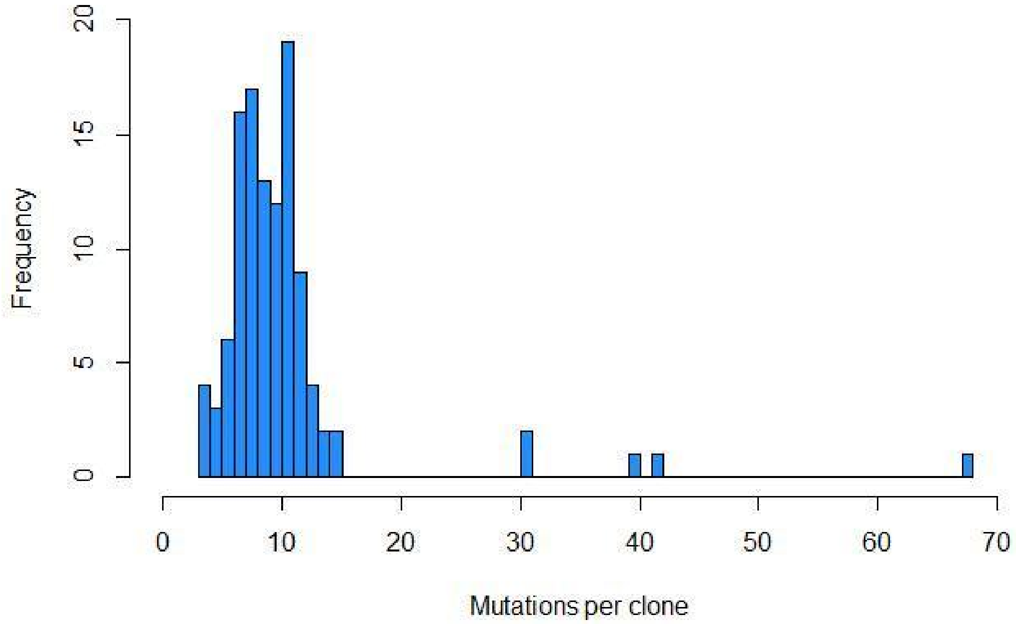


**Fig. S6:** Histogram of the number of mutations per clone, for all 112 populations.

### 5. Overview of genomics data

We first considered the number of mutations per mutation class: SNPs are the most common mutations, whereas Indels and SVs occur at lower rates (**Fig. S7**; see section 3 for the definition of mutation classes). The large and small populations evolved in increasing CTX concentrations have fixed more mutations per clone than the control populations with only tetracycline or no antibiotics (**Fig. S8**). When we consider the distribution of mutations in different classes for different treatments (**Fig. S9**), we see clear differences. Clones from large populations have accumulated more SNPs from small populations, whereas clones from large populations have fewer SVs than those from small populations. The tetracycline control has fixed equal numbers of each mutation class, whereas for the no antibiotics control, the pattern is more similar to the large populations: SNPs predominate but overall the numbers of mutations is lower. We can compare the mutation classes in large and small populations, considering whether mutations occur on the bacterial chromosome or the pACTEM plasmid (**Fig. S10**). The number of SNPs on the bacterial chromosome is similar for both treatments, whereas there are more SNPs on the plasmid in the large populations. A detailed description of the SVs on the bacterial chromosome is given in **Tables S4 and S5**. The number of SVs on the chromosome is higher for the small populations.

We also provide a comprehensive overview of mutations per clone from the evolved populations in **Fig. S11**, which includes the control populations.

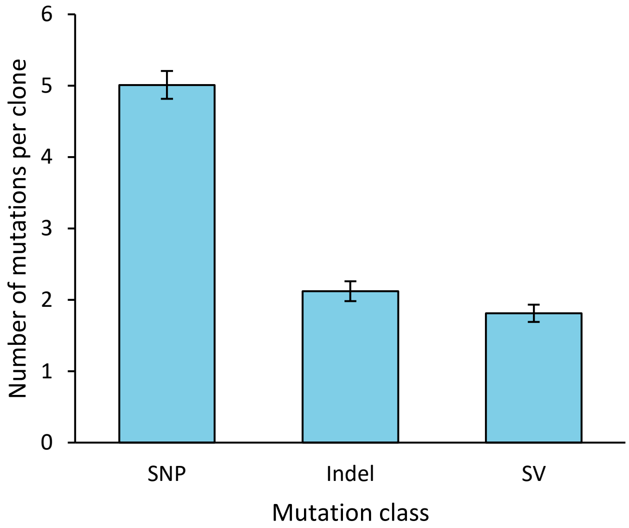


**Fig. S7:** Mutation frequency per mutation class for all 107 non-mutator sequenced clones from evolved populations, with error bars indicating the standard error of the mean. SNP: single nucleotide polymorphism; Indel: indels (<1 kbp) and IS-element insertions; SV: structural variants (> 1 kbp).

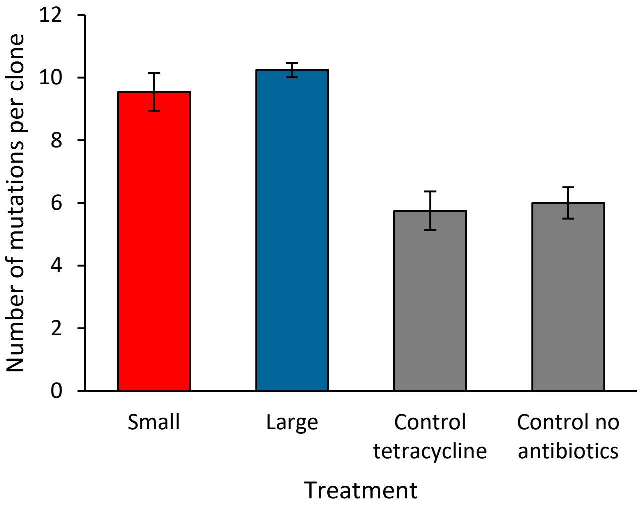


**Fig. S8:** Total mutation frequency per treatment for the 107 non-mutator populations, with error bars indicating the standard error of the mean.

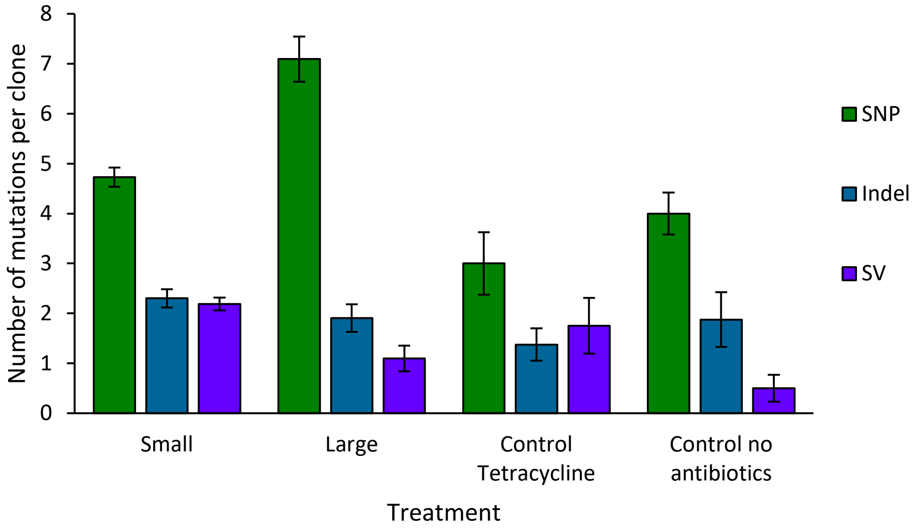


**Fig. S9:** Mutation frequency per mutation class and per treatment for the 107 non-mutator populations. Error bars indicate the standard error of the mean.

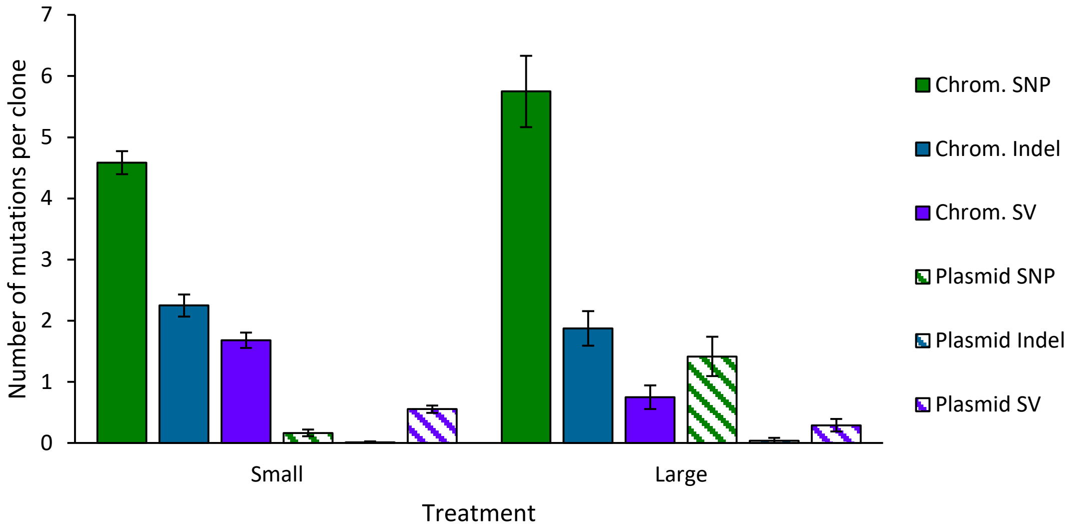


**Fig. S10:** Mutation frequency per mutation class in chromosome and plasmid, for large and small non-mutator populations only. Solid bars indicate the bacterial chromosome, whereas striped bars indicate the pACTEM plasmid. SNPs are green, Indels are blue, and SVs are purple. Error bars indicate the standard error of the mean.

**Fig. S11 (see next page)**: An overview of mutations across chromosome and plasmid per clone from the 112 evolved populations. Control populations are shown at the bottom of the figure, and small and large populations are shown in order of CTX resistance, as indicated to the left in MIC doublings. Small populations have a light blue filling, whereas large have none. The bacterial chromosome is indicated on the center left, and the pACTEM plasmid on the right. Chromosomal positions of rRNA operons and IS elements are shown above. Red and green bars indicate deletions and duplications, respectively, with a size over 1 kbp; yellow bars indicate deletions which have not fixed in the population, which were only observed in the plasmid; downward pointing green triangles and upward pointing red triangles indicate insertions and deletions under 1 kbp, respectively; black circles indicate IS insertions; orange diamonds indicate inversions; dark grey lines are intergenic or synonymous SNPs; black lines are nonsynonymous SNPs; asterisks indicate the five mutator clones.

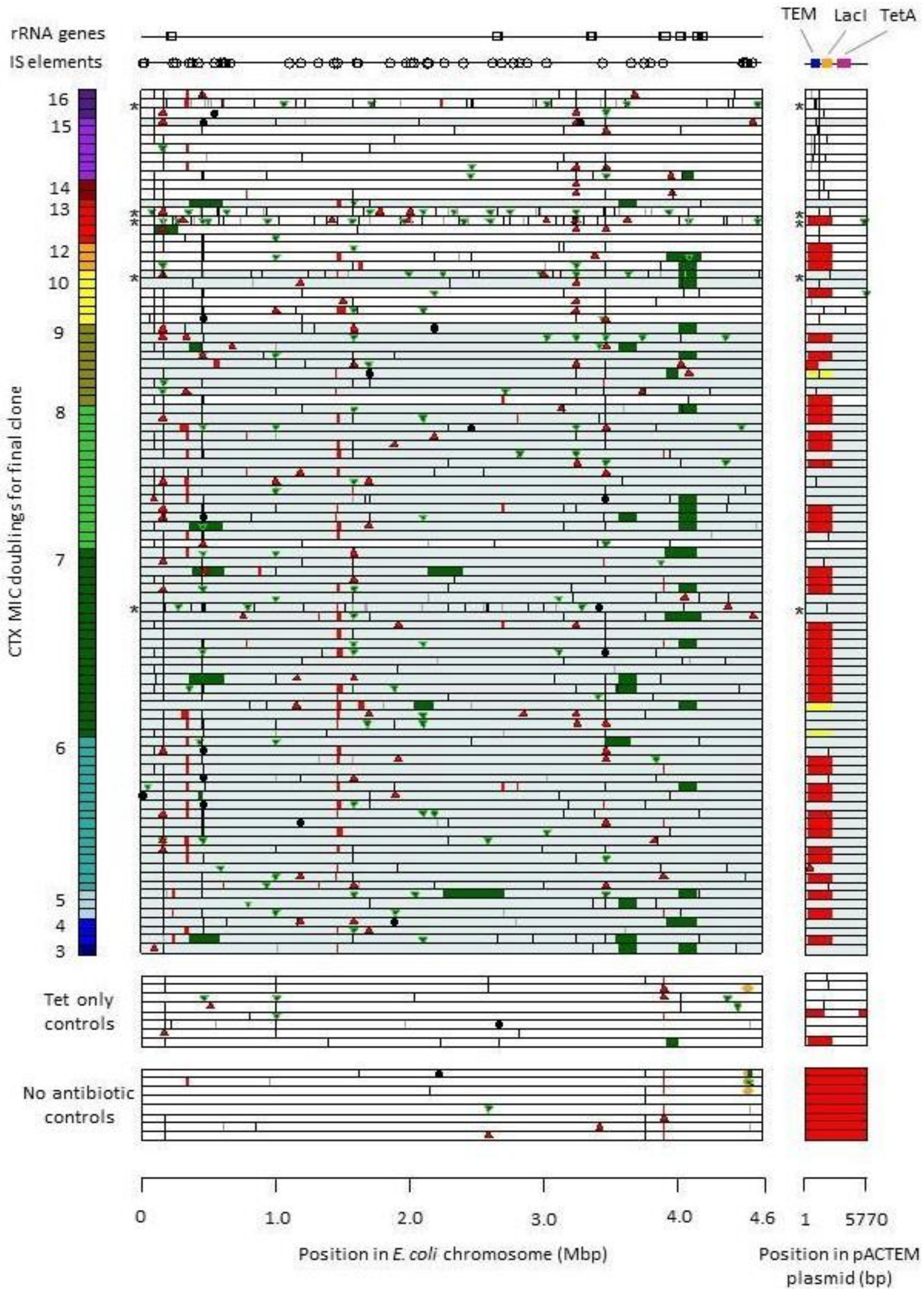


**Fig. S11:** Comprehensive overview of mutations for all 112 populations. The legend is given on the previous page.

**Table S4: Common deletions in evolved clones**

| Range[a] | Size[b] | Event[c] | Deleted genes | Occurrence | | | |
|---|---|---|---|---|---|---|---|
| | | | | S[d] | L[e] | N[f] | T[g] |
| 335,889-353,264 | 17,365 | IS-mediated | *frmAB, lacIZ, mhpABCDEFRT, yaiLO* | 20 | 4 | 1 | 0 |
| 1,460,931-1,466,876 | 5,946 | IS-mediated | *cybB, hokB, mokB, trg, ydcAIJ,* | 35 | 2 | 0 | 0 |
| 3,894,997-3,897,893 | 2,896 | IS150-mediated | *rbsACD* | 7 | 0 | 6 | 2 |

[a] The region affected that is shared by these deletions, [b] The size of she shared region denoted by range, [c] Type of event associated with the deletion. [d] Small populations, [e] Large populations, [f] No antibiotics control populations, [g] Tetracycline only control populations.

**Table S5: Common duplications in evolved clones**

| Range[a] | Size[b] | Duplicated genes | S[c] | L[d] | N[e] | T[f] |
|---|---|---|---|---|---|---|
| 429,491-457,290 | 28,429 | *acrAB, amtB, lon, cof, hha, hupB, glnK, mdlA, maa, ppiD, tesB, ybaABCEJOVWXYZ* | 7 | 0 | 0 | 0 |
| 3,651,100-3,652,225 | 1,125 | *insJ, insK-4* | 9 | 1 | 0 | 0 |
| 4,047,731-4,130,407[g] | 82,676 | *cdh, cpxAPR, csqR, cytR, dtd, eptC, fdhDE, fdoGHI, fief, fpr, frvABRX, frwBCD, fsaB, ftsN, gldA, glpFKX, hslUV, katG, kdgT, menA, metBFJL, pfkA, pflCD, priA, ptsA, rraA, rhaABDMSRT, rpmE, sbp, sodA, tpiA, uspD, yihRSTUVXY, yiiEFGMQRSX, yijEFO, zapB, ECB_03786, ECB_03822* | 19 | 3 | 0 | 1 |

[a] The region affected that is shared by these duplications, [b] The size of she shared region denoted by range, [c] Small populations, [d] Large populations, [e] No antibiotics control populations, [f] Tetracycline only control populations. [g] Many duplications in this region appear to depend on recombination between rRNA sequences.

### 6. dN/dS and dI/dS analysis

To determine the role of natural selection, we considered the ratio of the rate of nonsynonymous to the rate of synonymous substitutions (dN/dS). To generate an expectation of the number of nonsynonymous and synonymous mutations that could occur within the whole genome (bacterial chromosome and pACTEM plasmid), we assumed a plasmid copy number of 10, and included the effect of mutational bias based on mutation accumulation data for *E. coli* [18]. We then determined the rate for each mutation class, by dividing the observed mutations by the expectation. We also considered the ratio of the rate of intergenic mutations to the rate of synonymous substitutions (dI/dS), as previously suggested [19], to get an indication of whether mutations in promotor regions were overrepresented in the data. To estimate the percentage of beneficial mutations, we assumed that all nonsynonymous or intergenic mutations that occur above the baseline substitution rate (as determined by the synonymous mutations) were beneficial, e.g. percentage beneficial mutations = 100*(dN/dS -1)/(dN/dS).

We generally found dN/dS and dI/dS values well above 1 for the non-mutator clones (**Table S6**). By contrast, for the mutator clones dN/dS ~1, as expected for populations substituting mutations by chance. Large populations had appreciably higher dN/dS and dI/dS values than the other treatments, indicating stronger effects of selection in these populations.

**Table S6:** Results of dN/dS and dI/dS analyses**.**

| Clones | Populations | dN/dS | N beneficial | dI/dS | I beneficial |
|---|---|---|---|---|---|
| Mutators | All | 1.302 | 23.2% | 0.733 | - |
| Non-mutators | All | 13.789 | 92.7% | 4.692 | 78.7% |
| | Small | 11.938 | 91.6% | 3.910 | 74.4% |
| | Large | 28.830 | 96.5% | 7.819 | 87.2% |
| | No antibiotic | 6.234 | 84.2% | 5.864 | 82.9% |
| | Tetracycline only | 11.532 | 91.3% | 5.864 | 82.9% |

### 7. Repeatability of genomic changes

#### *Methodology*

We wanted to quantify how repeatable genome evolution was with metrics that could compare mutational events of all sizes, based on either (i) the exact position of mutations, or (ii) based on the genes modified by these mutations. We therefore formulated the *H*-index of repeatability, which for the pairwise comparison of a genotype *A* to *B* with *m* and *n* mutations, respectively, is:

$$H_{A,B} = \sum_{j=1}^{m}\sum_{k=1}^{n}\left(\frac{A_j B_k}{A_j}\right) \Big/ \sum_{j=1}^{m}\sum_{l=1}^{m}\left(\frac{A_j A_l}{A_j}\right) \quad (1)$$

The numerator measures the similarity between two genotypes and the denominator normalizes by the similarity of a genotype to itself, as mutational events can overlap. E.g., a SNP can occur in a region that is duplicated in another genotype. As this metric is asymmetrical, the repeatability measure for the two genotypes is the mean of both reciprocal comparisons:

$$H = \frac{H_{A,B} + H_{B,A}}{2} \quad (2)$$

If the metric is performed with the exact coordinates of mutations, it is a nucleotide-level *H*-index, whereas if the genes affected by mutations are used, it is the gene-level *H*-index. When *H* is zero, there is no repeated evolution, whereas values of *H* of 1 represent convergence on the same genotype. For determining the *H*-index for *N* genotypes, we first determined the mean *H*-index for each genotype versus all other genotypes, and then performed further analyses on these data (as opposed to analyzing all pairwise *H*-indexes), avoiding inflation of the degrees of freedom.

#### *Nucleotide-level repeatability results*

We determined nucleotide level *H*-indexes for clones from all evolved non-mutator populations, considering different classes of mutations. First, if all mutations are combined then *H*-indexes are lower for the small than for the large populations (**Fig. S12A**). If we consider the *H*-index for the three different mutation classes (SNPs, Indels and SVs), it is apparent that different mutation classes drive overall repeatability of the small and large populations. In clones from the small populations, SVs show higher convergence than in clones from the large populations (**Fig, S12B**). The control without antibiotics has a high overall *H*-index (**Fig. S12A**), which is driven by SVs (**Fig. S12B**), particularly the loss of the pACTEM plasmid in all lineages (**Fig. S11**). The tetracycline only control has a more similar profile to the large populations, with SNPs being more convergent than Indels or SVs (**Fig. S12**). We also calculate *H*-indexes separately for the genome and plasmid (**Table S7**). This analysis demonstrates that the convergent evolution seen for SNPs in the large populations occurs on both plasmid and bacterial chromosome, so it is not driven only by recurring mutations affecting the TEM-1 gene on the pACTEM plasmid.

Tenaillon et al. [19] report another statistic for quantifying the repeatability of molecular evolution, in which events that overlap for more than 10% of their length are considered to be repeated events, which we refer to as the *T*-index. The *H*-index and *T*-index are both asymmetrical and require reciprocal comparisons to be made between each pair of genotypes, but a possible advantage of the *H*-index is that it does not require the arbitrary choice of threshold value for overlap between events. We compared the two indexes for our data and found that they generally give very similar results (**Fig. S13**).

#### *Gene-level repeatability results*

We determined gene-level *H*-indexes for clones from all evolved non-mutator populations, considering different classes of mutations. As expected, gene-level repeatability was higher than nucleotide-level repeatability (**Fig. S14**). One striking difference between the nucleotide-level and gene-level *H*-indexes is for the small populations, where repeatability for SNPs is much higher at the gene-level. Hence, although SNPs are less likely to affect the same nucleotide positions in small populations, they are likely to affect the same genes. As with the nucleotide-level *H*-indexes, the no antibiotics treatment has a higher repeatability than the tetracycline only treatment, although the no antibiotics treatment also has convergence for SNPs at the gene level (**Figs. S12 and S14**). All calculated gene-level *H*-indexes are given in **Table S8**.

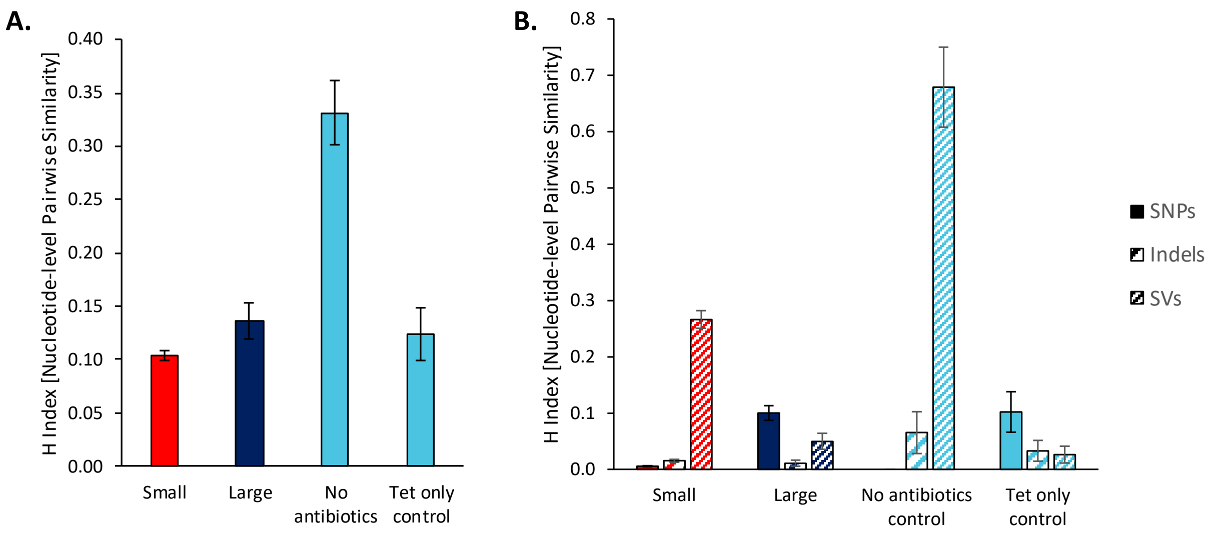


**Fig. S12:** (**A**) Nucleotide-level *H*-index for all mutational events in clones from non-mutator populations. (**B**) Nucleotide-level *H*-index for three classes of mutational events, as indicated by the legend.

**Table S7:** Nucleotide-level *H*-indexes for different loci (bacterial chromosome and the pACTEM plasmid) and different classes of mutations in clones from non-mutator populations. Mean *H*-index values ± standard error of the mean are given.

| | | Treatment | | | |
|---|---|---|---|---|---|
| Locus | Mutations | Small | Large | No antibiotics | Tetracycline |
| Both | All | 0.104 ±0.005 | 0.136 ±0.075 | 0.331 ±0.030 | 0.124 ±0.025 |
| Chromosome | All | 0.068 ±0.003 | 0.070 ±0.011 | 0.176 ±0.032 | 0.116 ±0.024 |
| Plasmid | All | 0.361 ±0.029 | 0.352 ±0.043 | 1.000 ±0 | 0.132 ±0.048 |
| Both | SNP | 0.006 ±0.001 | 0.100 ±0.060 | 0 ±0 | 0.102 ±0.036 |
| | Indel | 0.015 ±0.002 | 0.011 ±0.005 | 0.066 ±0.037 | 0.033 ±0.019 |
| | SV | 0.266 ±0.133 | 0.050 ±0.014 | 0.679 ±0.071 | 0.026 ±0.015 |
| Chromosome | SNP | 0.006 ±0.001 | 0.070 ±0.016 | 0 ±0 | 0.118 ±0.040 |
| | Indel | 0.015 ±0.002 | 0.011 ±0.005 | 0.066 ±0.037 | 0.033 ±0.019 |
| | SV | 0.179 ±0.014 | 0.021 ±0.008 | 0.336 ±0.091 | 0.030 ±0.020 |
| Plasmid | SNP | 0.001 ±0.001 | 0.136 ±0.034 | 0 ±0 | 0 ±0 |
| | Indel | 0 ±0 | 0 ±0 | 0 ±0 | 0 ±0 |
| | SV | 0.285 ±0.032 | 0.048 ±0.019 | 1.000 ±0 | 0.010 ±0.007 |

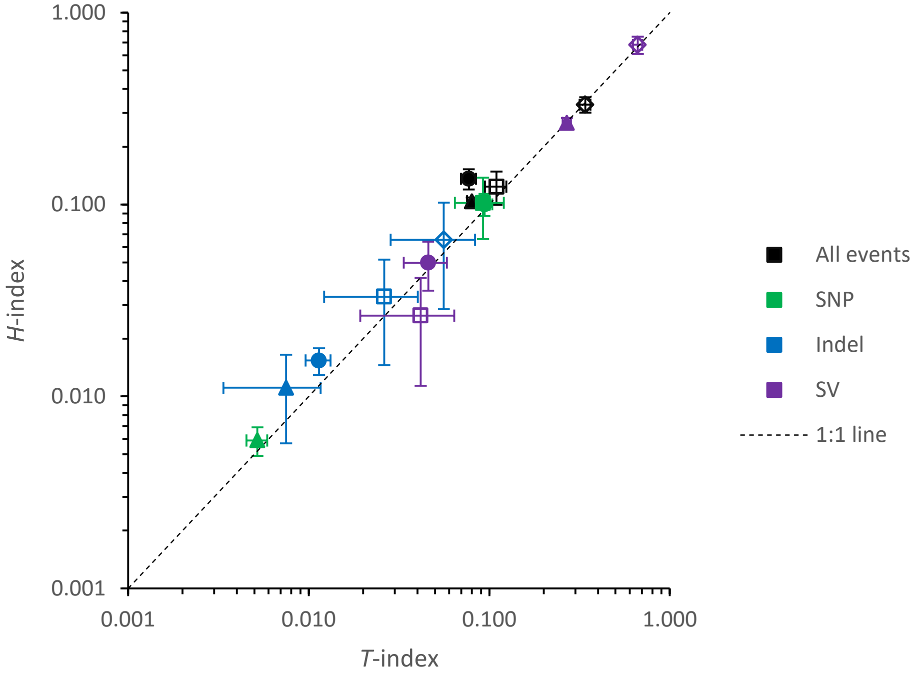


**Fig. S13:** Comparison of the nucleotide-level *T* [19] and *H*-indexes for the final clone data. Each data point represents the mean *H*-index value for the small (filled triangles), large (filled circles), no antibiotics (open diamonds) or tetracycline only (open squares) treatments, for either all mutations (black), SNPs (green), Indels (blue) or SVs (purple), and error bars indicate the standard error of the mean. The 1:1 line is given as a reference only. For all the square-root-transformed mean *T* and *H*-indexes included in the figure, the coefficient of determination ($r^2$) is 0.983, indicating good agreement between the different indexes for these data.

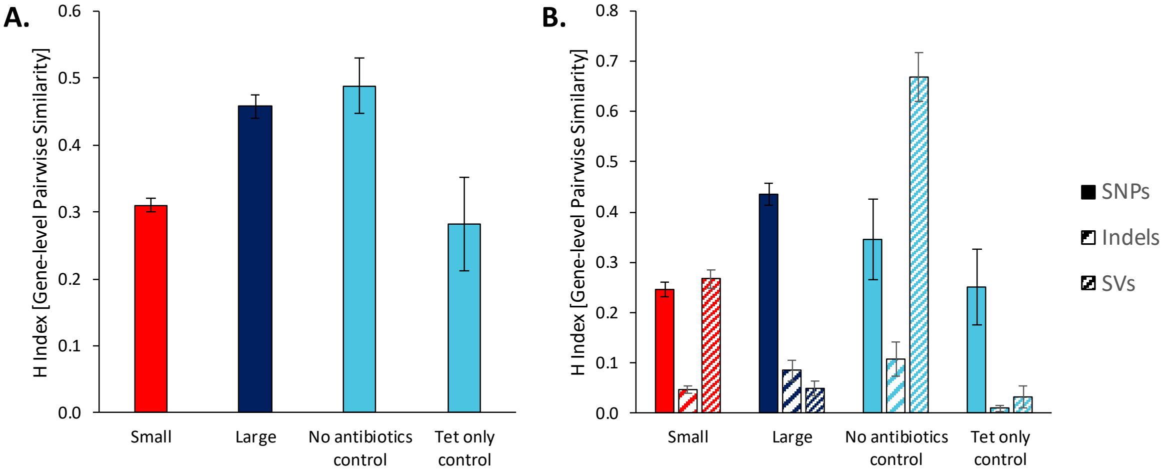


**Fig. S14:** (**A**) Gene-level *H*-index is given for all mutational events in clones from non-mutator populations. (**B**) Gene-level *H*-index is given for three classes of mutational events, as indicated by the legend.

**Table S8:** Gene-level *H*-indexes for different loci (bacterial chromosome and the pACTEM plasmid) and different classes of mutations in clones from non-mutator populations. Mean *H*-index values ± standard error of the mean are given.

| | | Treatment | | | |
|---|---|---|---|---|---|
| Locus | Mutations | Small | Large | No antibiotics | Tetracycline |
| Both | All | 0.310 ±0.010 | 0.458 ±0.018 | 0.488 ±0.042 | 0.281 ±0.070 |
| Chromosome | All | 0.284 ±0.010 | 0.407 ±0.039 | 0.358 ±0.052 | 0.249 ±0.064 |
| Plasmid | All | 0.448 ±0.038 | 0.831 ±0.125 | 1.000 ±0 | 0.321 ±0.100 |
| Both | SNP | 0.246 ±0.014 | 0.435 ±0.021 | 0.345 ±0.080 | 0.251 ±0.076 |
| | Indel | 0.046 ±0.007 | 0.084 ±0.021 | 0.107 ±0.033 | 0.009 ±0.006 |
| | SV | 0.267 ±0.018 | 0.049 ±0.015 | 0.668 ±0.050 | 0.032 ±0.021 |
| Chromosome | SNP | 0.251 ±0.015 | 0.386 ±0.050 | 0.345 ±0.080 | 0.231 ±0.068 |
| | Indel | 0.046 ±0.007 | 0.084 ±0.021 | 0.107 ±0.033 | 0.009 ±0.006 |
| | SV | 0.176 ±0.015 | 0.020 ±0.007 | 0.314 ±0.072 | 0.032 ±0.021 |
| Plasmid | SNP | 0.009 ±0.003 | 0.401 ±0.084 | 0 ±0 | 0.214 ±0.081 |
| | Indel | 0 ±0 | 0 ±0 | 0 ±0 | 0 ±0 |
| | SV | 0.287 ±0.032 | 0.048 ±0.019 | 1.000 ±0 | 0 ±0 |

#### 8. Regression analysis of the relative frequency of SVs

To investigate whether the differences in selective conditions between small and large populations could explain the mutational repeatability pattern of SNPs and SVs, we performed linear regression analyses of the relationship between the fraction of SVs with respect to all mutations and the geometric mean of the CTX concentrations during the evolution experiment. Indeed, across all populations a significant dependence of the fraction of SVs on selective CTX concentrations is apparent (**Fig. S15**, $R^2$=0.31, $P$<0.001). However, this dependence is not even approaching significance when considering either only small or large populations (Small; $R^2$=0.01, $P$=0.34, Large; $R^2$=0.02, $P$=0.59). Note that the lack of effect from selective CTX concentration among populations of the same size is not due to lack of variation in selective CTX concentration. This suggests that differences in mutation supply, and not differences in selective conditions, are responsible for the increased fraction of SVs in small populations. The regression analyses were performed using R version 3.6.1.

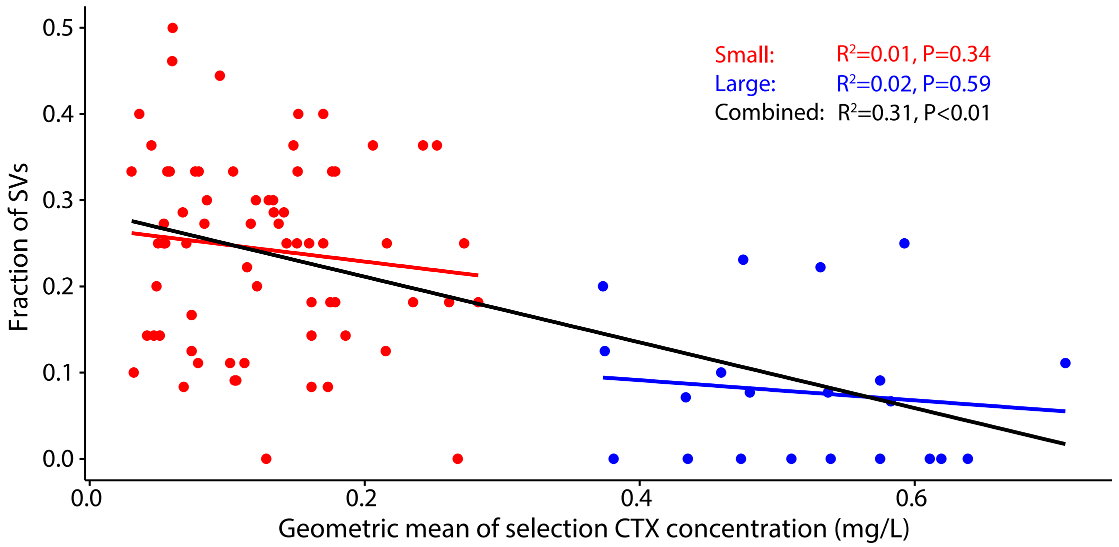


**Fig. S15**. Regression analysis of the fraction of SVs among all mutations per clone against CTX concentration.

### 9. Dynamics of genomic changes

#### *Muller plots*

Population samples from different time points were analyzed with a metagenomics approach (see also "Detection of mutations for population samples" in section 2 of this document). A single clone from the final time point was also sequenced for each of these populations (see Section 4 "Overview of Genomics Data" of this document), and we checked whether the two sequencing results were congruent. If the final populations are polymorphic, we expect to detect some mutations only in the population and not all mutations detected in the population in the clone, which is congruent with our results for populations L1, L2, L5 and S4 (**Table S9**). We expect that a small number of mutations may be detected in the clones but not in the populations because they were present at low frequency, and we indeed found a total of eight mutations only in the clones (**Table S9**). By contrast, we expect to find mutations that are fixed in the population in the clones, which we found for 83 out of 85 cases (**Table S9**). For the two mutations that were fixed in the meta-populations but not found in the clones, there was a reasonable explanation for why these discrepancies occurred (see footnotes in **Table S9**). The data for the clones and populations are therefore generally in good agreement.

Using the combined clone and metapopulation sequencing results, we were able to determine the main haplotypes in the populations heuristically. For some of the small populations this was straightforward, as there was little clonal interference and most mutations detected fix successively along the single line of descent (S1, S14, S17 and S25). In other cases, the dynamics were more complex. The final clone data helped to identify the variants that coexisted for at least 200 generations in S4, and had similar frequencies at the time of measurement. In the large populations, at some time points there were multiple mutations that became extinct, and we could not establish whether there was linkage between them. For example, the multiple non-line-of-descent mutations in *acrB* appeared to occur on different read pairs, suggesting they were two different haplotypes, but this was an exceptional case. In other cases, mutations were assumed to be linked depending on the correlation of their frequencies. We used the *R* library *fishplot* [20] to visualize these data (**Fig. 4A**).

#### *Repeatability of evolution*

To determine whether the repeatability of the metagenomics results was similar to that obtained for the clones, we also estimated *H*-indexes for the time course data. We first performed this analysis for the majority mutations (frequency > 0.5) at the final time point (**Fig. S16A**). This gave similar patterns for small and large populations to those observed for the clones (see Section 5 "Repeatability of genomic changes"), albeit with more uncertainty due to the much smaller number of replicates. In the small populations, there was a high frequency of convergent SVs, whereas there were none in the large populations (**Fig. S16A**). SNPs showed a higher repeatability in large than in small populations, although there was some convergence in the small populations driven by the occurrence of TEM mutations R241P in two of the small populations.

We also considered repeatability for all mutations detected – at any frequency – in the metagenomes over all time points, irrespective of their frequency, and hence fate (extinction or fixation). Here we saw patterns that were less divergent for the small and large populations, and strikingly SV repeatability was identical for both population sizes (**Fig. S16B**). This finding suggests that the mutation supply of SV is similar in large and small populations, but that clonal interference with large-effect point mutations leads to the loss of beneficial SVs in large populations, in support of our hypothesis.

**Table S9**: Comparison of mutations detected in clones and final populations. The number of mutations is given, with the percentage of all detected mutations per clone-population pair shown in parentheses.

| | Unique mutations[a] | | | Shared mutations[b] | |
|---|---|---|---|---|---|
| | Clone | Population | | | |
| Population | | Polymorphic | Fixed | Polymorphic | Fixed |
| L1 | 2 (14.3%) | 5 (35.7%) | 0 (0%) | 0 (0%) | 7 (50.0%) |
| L2 | 1 (6.7%) | 4 (26.6%) | 0 (0%) | 3 (20.0%) | 7 (46.7%) |
| L3 | 1 (8.3%) | 0 (0%) | 0 (0%) | 0 (0%) | 11 (91.7%) |
| L4 | 0 (0%) | 0 (0%) | 0 (0%) | 0 (0%) | 15 (100%) |
| L5 | 1 (12.5%) | 1 (12.5%) | 1 (12.5%)[c] | 1 (12.5%) | 4 (50.0%) |
| S1 | 0 (0%) | 0 (0%) | 1 (7.7%)[d] | 1 (7.7%) | 11 (84.6%) |
| S4 | 0 (0%) | 3 (16.7%) | 0 (0%) | 10 (55.6%) | 5 (27.8%) |
| S14 | 0 (0%) | 0 (0%) | 0 (0%) | 0 (0%) | 11 (100%) |
| S17 | 1 (11.1%) | 0 (0%) | 0 (0%) | 0 (0%) | 8 (88.9%) |
| S25 | 2 (22.2%) | 0 (0%) | 0 (0%) | 0 (0%) | 7 (77.8%) |
| Sum | 8 (6.5) | 14 (11.3) | 2 (1.6) | 17 (13.7) | 83 (66.9) |

[a] Mutations found only in the sequenced final clone or the meta-population. [b] Mutations found in both the sequenced clone and meta-population. [c] The fixed mutation from the meta-populations that was not detected in the clone (10 base-pair deletion in *nlpI*), was present a frequency of 0.688 at 400 generations

and 1.000 at 500 generations. Although our meta-population sequencing suggested that the mutation had gone to fixation (depth was only ≈ 24 at this site), in reality this mutation was at high frequency but not fixed and the clone represents a minority variant. [d] The duplication 4,013,902-4,146,518 was detected in the clone, but its frequency was too low to be considered reliable (1.3 fold increase in coverage). Given that this duplication was present in the meta-population, presumably a reversion of this duplication occurred during the culturing of the clone.

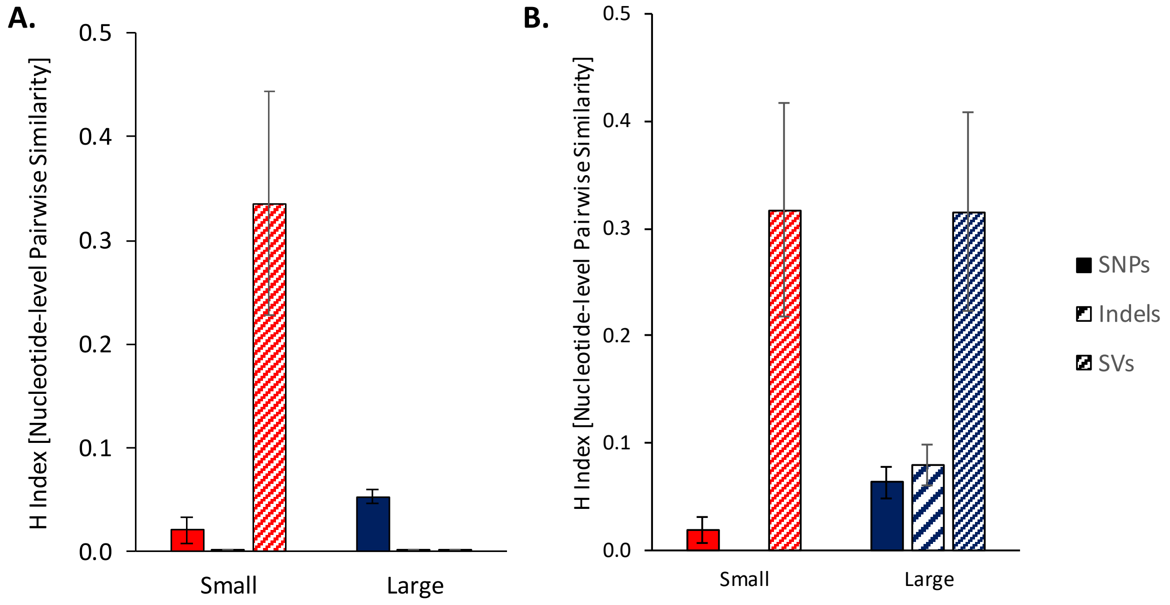


**Fig. S16.** Nucleotide-level *H*-indexes for the five small and five large populations of **Fig. 4A** for which we obtained time-course meta-genome data, with fill pattern indicating the mutation category. (**A**) *H*-index for the majority mutations (frequency > 0.5) at the final time point. (**B**) *H*-index for all mutations detected at all timepoints, regardless of their frequency. Since the majority of detected mutations reach a majority frequency in the small populations, the indexes for detected and majority mutations are very similar. In the large populations, convergent SVs are detected, but clonal interference with large-effect SNPs prevents these mutations from fixing.

### 10. Inference of mutation parameters from Wright-Fisher simulations

We consider standard asexual Wright-Fisher (WF) dynamics [3] of a population where individuals can acquire beneficial mutations from three different classes. Mutations in each class occur randomly and independently with rates $U_i$ per class $i = 1,2,3,$ individual and generation. The numbers of possible mutations for each class are given by $(L_1, L_2, L_3) = (20,6,6)$, which is sufficiently large such that the mutation supplies are not depleted at the end of the simulation for the parameter ranges expected to fit the experimental data. Thus the mutation rates for each single mutation are given by $U_i/L_i$. The genotypes $\tau$ of individuals in this model are encoded by three binary sequences $\tau_i$ of length $L_i$ where each element is 1 or 0 indicating the presence or absence of the corresponding mutation. The selection coefficient of each mutation is chosen randomly from an exponential distribution with mean $s_i$ for mutation class $i$. A mutation with selection coefficient $s$ increases the fitness $f$ of the individual by a factor $1 + s$, and the effects of different mutations combine multiplicatively (no epistasis).

The WF dynamics is implemented as follows. At each time step $t$, the expected number of individuals of genotype $\tau$ at time $t + 1$ without selection is computed as:

$$P_{t+1}(\tau) = \left(1 - \sum_{i=1}^{3} U_i\right) N_t(\tau) + \sum_{i=1}^{3} \frac{U_i}{L_i} \sum_{j=1}^{L_i} N_t(\Delta_j^i \tau), \qquad (7)$$

where $N_t(\tau)$ is the number of individuals of genotype $\tau$ in generation $t$ and $\Delta_j^i \tau$ stands for the genotype obtained by flipping the $j$-th binary variable of $\tau_i$. In the selection step, these numbers are then normalized with weights given by their fitness values $f(\tau)$,

$$\tilde{P}_{t+1}(\tau) = \frac{P_{t+1}(\tau) f(\tau)}{\sum_{\tau'} P_{t+1}(\tau') f(\tau')}. \qquad (8)$$

The genotypes are finally sampled $N$ times independently with a probability proportional to $\tilde{P}_{t+1}(\tau)$ to form a final population of total size $N$. The details of the algorithm are described in [21].

Following the setting of the experiment, we carried out WF simulations up to $t = 500$ generations for two different population sizes $N = 2 \times 10^6$ and $N = 2 \times 10^8$. Initially, the entire population is monomorphic for the wildtype genotype with fitness $f = 1$. Our aim is to determine the six model parameters $\{U_i, s_i\}_{i=1,2,3}$ such that the numbers of fixed mutations $C_i$ for each class and each population size at the end point of the experiment are reproduced. This requires in principle to obtain the full distribution of $C = \{C_i^{small,large}\}_{i=1,2,3}$ and perform a likelihood maximization. Unfortunately, since the statistics of $C$ are accessible only through explicit simulations, this is a computationally daunting task that is not feasible in practice.

Instead, we approximate this distribution using a machine learning approach. Specifically, we use a so-called mixture density network with a Gaussian ansatz [22, 23]. We train the neural network to establish a functional relation:

$$g_w : \{U_i, s_i\}_{i=1,2,3} \to \{m_i^{small}, m_i^{large}, \sigma_i^{small}, \sigma_i^{large}\}_{i=1,2,3}, \quad (9)$$

where $m_i$ and $\sigma_i$ is the mean and the standard deviation of the random variable $C_i$. Correlations between the $C_i$ values of different classes are neglected. The training procedure is implemented as follows. For an initially arbitrarily chosen parameter vector, which we call the center $\mathcal{P} = \{\ln U_i, s_i\}_{i=1,2,3}$, we create a cloud of parameter vectors $\mathcal{P}_i = \mathcal{P} + \xi_i$ of size 100, where the $\xi_i$ values are randomly generated vectors. Then, for each parameter vector $\mathcal{P}_i$, we run two WF simulations to obtain $C_i^{\text{small}}$ and $C_i^{\text{large}}$ for the two different population sizes. This parameter-result pair is then added to the training set. Once this step is finished for the entire cloud, we train the neural network by maximizing

$$\mathcal{L} = \sum_i \left( \ln \text{Prob}^{\text{small}}\left(C_i^{\text{small}} \middle| \mathcal{P}_i\right) + \ln \text{Prob}^{\text{large}}\left(C_i^{\text{large}} \middle| \mathcal{P}_i\right)\right) \quad (10)$$

over the weights $w$, where $\text{Prob}(C_i|P_i)$ is a Gaussian distribution with parameters estimated from the network $g_w$.

After the training step, the next center is chosen to be the parameter vector with the largest log-likelihood for the empirical data. The whole process is repeated until the position of the center converges to a fixed value. By construction, this process only increases the size of training set (by 100 per iteration), and once the size of the training set is sufficiently large, it is expected that the neural network fully learns the distribution at least around the maximum.

In our actual implementation, we choose to approximate $g_w$ as a fully connected feed-forward neural network with one hidden layer of 100 hidden units. The activation functions for these hidden neurons as well as the output neurons for the mean values $m_i$ are chosen to be ReLU (rectified linear unit) activation functions. Since the standard deviations $\sigma_i$ are positive by definition, this condition is imposed through exponential linear units. The random variables $\xi_i$ for cloud generation are chosen to be Gaussian random vectors with standard deviation 0.1. The final size of the training set is reached at $10^5$. We tested our approach by choosing different initial centers and found that they converged to the same final point. Finally, we performed a mini-batch optimization with Adam optimizer with an initial rate of $10^{-3}$. The inferred model parameters are summarized in **Table S10**, and **Fig. S17** compares the distribution of mutation numbers obtained from the WF model with optimized parameters to the experimental data.

**Table S10:** Inferred mutation rates and selection coefficients for the three mutation classes.

| | SNP | Indel | SV |
|---|---|---|---|
| $U_i$ | $2.17 \times 10^{-8}$ | $1.763 \times 10^{-7}$ | $7.051 \times 10^{-6}$ |
| $s_i$ | 0.413 | 0.250 | 0.138 |

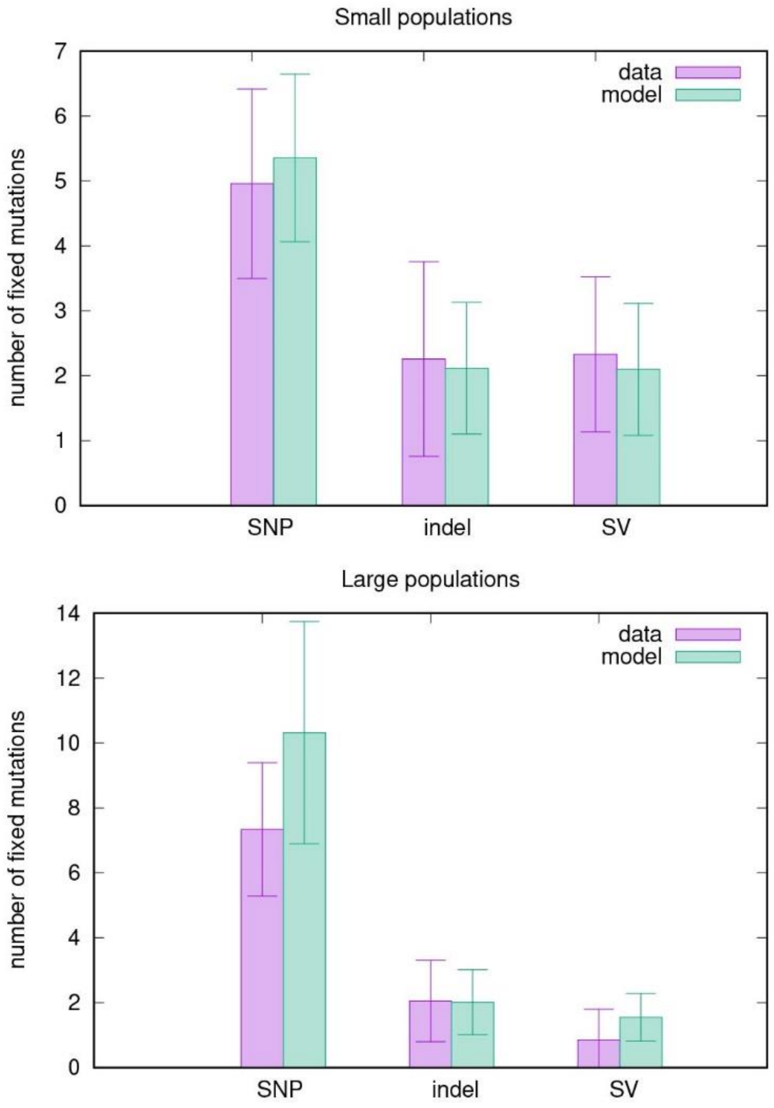


**Fig. S17.** Distribution of fixed mutations at the evolutionary endpoint obtained from the optimized WF model in comparison to the experimental data. Column heights represent the mean number of mutations and error bars show the corresponding standard deviations.

### 11. MIC effects of different mutation classes

For the clones from the evolved populations, we estimated MIC-effect sizes for different classes of mutations, using the genome data and CTX resistance data. To predict the effect size, we fit a series of general linear models to the data that predict the increase in resistance, $\Delta R$, such that $\Delta R = log_2(MIC_{evolvedclone}) - log_2(MIC_{ancestor})$. There are 12 types of mutations possible (3 classes [SNP, Indel and SV] × 2 loci [chromosome and plasmid] × 2 population types [large and small]), so maximally 12 coefficients need to be estimated (**Fig. S18**). In the simplest case, we assume all mutations have the same effect on MIC (Model 1), and therefore $\Delta R = \lambda q$ where $\lambda$ is the effect-size coefficient (to be estimated from the data) and $q$ is the total number of mutations of all types. Note there is no intercept in any of the models. Model 2 assumes mutations on the plasmid and chromosome – denoted by subscripts $p$ and $c$ – can have different effect sizes and hence $\Delta R = \lambda_c q_c + \lambda_p q_p$, and so forth for all models including the full model with 12 coefficients. All eight models were fitted to the data in R assuming a Normal distribution, and model selection was performed with the Akaike information criterion (AIC).

Model selection results are reported in **Table S11**, and model parameter estimates are given in **Table S12**. Model selection suggests that Models 4, 6 and 8 are the best-supported models. Model 8 appears to be over-parametrized, given it is the most complex and least well supported of the three models. Model 6 (which assumes differences between mutation effect size in large and small populations, on the bacterial chromosome and plasmid) essentially predicts that mutational effect size is larger for mutations in the plasmid than in the bacterial chromosome, and that mutations in plasmids from large populations have larger effect sizes than in plasmids from small populations. The model estimates therefore are being driven by TEM mutations and whilst highlighting their importance, they do not shed light on effect sizes for different classes of mutations. Model 4 (three mutation classes, separately for chromosome and plasmid) has a similar fit and level of support to Model 6, but includes mutation classes instead of population size, and is therefore the most informative model on the differences between effect size for different classes of mutations.

For model 4, SNPs have a highly significant effect on resistance (**Table S12**), whereas the effect of SVs is marginally significant for the plasmid and insignificant for the bacterial chromosome. For events on the same locus, SNPs have a significant larger effect than SVs ($t$ test; chromosome: $t = 2.094$, d.f. = 180, $P = 0.038$, plasmid: $t = 3.455$, d.f. = 180, $P < 0.001$). The effect of indels is intermediate, but estimates are poor for the plasmid due to the low number of indels that occur here. By contrast, the number of Indels on the chromosome is larger, but the effect size is not significantly different from either SNPs ($t = 0.692$, d.f. = 180, $P = 0.490$) or SVs ($t = 1.416$, d.f. = 180, $P = 0.158$).

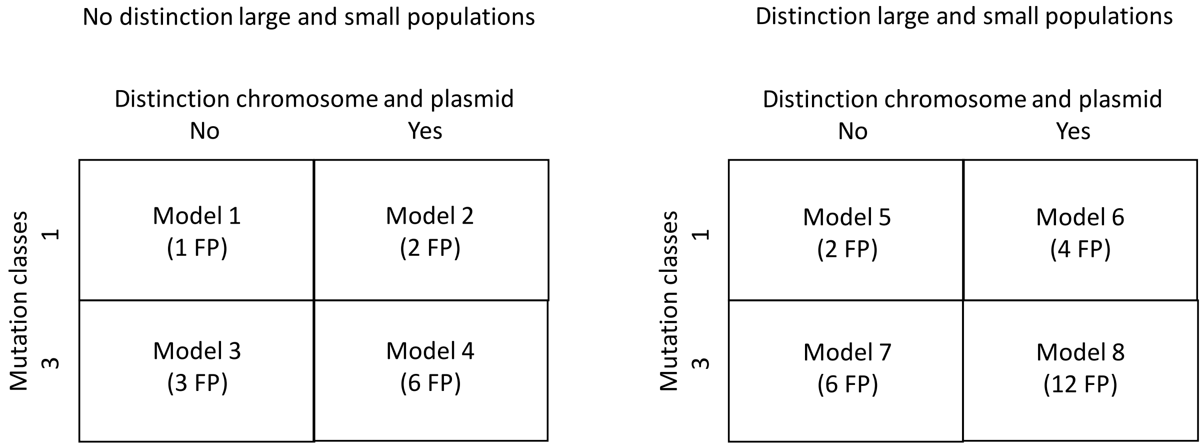


**Fig. S18:** Overview of the eight different mutational effect-size models fitted to the MIC data. FP stand for "free parameters". The three mutation classes used are SNP, Indel and SV.

**Table S11:** Model selection with the AIC for effect sizes of mutation classes.

| Model | Parameters | NLL | AIC | ΔAIC |
|---|---|---|---|---|
| 1 | 1 | 240.239 | 484.48 | 70.90 |
| 2 | 2 | 212.816 | 431.63 | 18.05 |
| 3 | 3 | 228.313 | 464.63 | 51.05 |
| 4 | 6 | 201.668 | 417.34 | 3.76 |
| 5 | 2 | 212.239 | 432.48 | 18.90 |
| 6 | 4 | 200.790 | 413.58 | - |
| 7 | 6 | 209.045 | 434.09 | 20.51 |
| 8 | 12 | 195.460 | 418.92 | 5.34 |

**Table S12:** Estimated model parameters and their standard errors of the mean (SEM). Subscripts c and p denote the chromosome or plasmid, and subscripts *snp*, *ind* and *sv* refer to SNP, Indels and IS-element insertions and SVs, respectively. For models 5-8 the parameter estimates are made separately for large and small populations. Significance levels for one-sample *t* tests against a value of zero are indicated by asterisks: *** $P$ < 0.001, ** $P$ < 0.01, * $P$ < 0.05.

| Model | Parameter estimate ±SEM |
|---|---|
| 1 | $\lambda$ = 0.835 ±0.035*** |
| 2 | $\lambda_c$ = 0.589 ±0.039***, $\lambda_p$ = 3.087 ±0.263*** |
| 3 | $\lambda_{snp}$ = 1.230 ±0.096***, $\lambda_{ind}$ = 0.717 ± 0.189***, $\lambda_{sv}$ = -0.132 ±0.204 |
| 4 | $\lambda_{c,snp}$ = 0.803 ±0.094***, $\lambda_{c,ind}$ = 0.684 ±0.144***, $\lambda_{c,sv}$ = 0.327 ±0.207,<br>$\lambda_{p,snp}$ = 3.180 ±0.250***, $\lambda_{p,ind}$ = 0.530 ±1.707, $\lambda_{p,sv}$ = 1.193 ±0.518* |
| 5 | Large: $\lambda$ =1.136 ±0.099***; Small $\lambda$ = 0.729 ±0.025*** |
| 6 | Large: $\lambda_c$ = 0.709 ±0.110***, $\lambda_p$ = 3.255 ±0.435***;<br>Small: $\lambda_c$ = 0.651 ±0.041***, $\lambda_p$ = 1.702 ±0.408*** |
| 7 | Large: $\lambda_{snp}$ = 1.310 ±0.252***, $\lambda_{ind}$ = 1.140 ±0.788, $\lambda_{sv}$ = -0.235 ±1.156;<br>Small: $\lambda_{snp}$ = 0.890 ±0.102***, $\lambda_{ind}$ = 0.759 ±0.140***, $\lambda_{sv}$ = 0.355 ±0.175* |
| 8 | Large: $\lambda_{c,snp}$ = 0.691 ±0.274*, $\lambda_{c,ind}$ = 1.126 ±0.638, $\lambda_{c,sv}$ = 0.283 ±1.321,<br>$\lambda_{p,snp}$ = 3.208 ±0.481***, $\lambda_{p,ind}$ = -0.684 ±4.103, $\lambda_{p,sv}$ = 2.652 ±2.850;<br>Small: $\lambda_{c,snp}$ = 0.775 ±0.107***, $\lambda_{c,ind}$ = 0.690 ±0.138***, $\lambda_{c,sv}$ = 0.387 ±0.205,<br>$\lambda_{p,snp}$ = 2.392 ±0.560***, $\lambda_{p,ind}$ = 1.048 ±2.006, $\lambda_{p,sv}$ = 1.110 ±0.480* |

#### 12. Analysis of functional targets and evolutionary trajectories

We defined functional targets, and their likely adaptive role in our evolution experiment, by grouping genes affected by at least five SNPs and/or indels in the 96 CTX-treated populations, based on information about their function and regulation in EcoCyc (ecocyc.org) and literature (references inserted below), as well as the types of mutations involved (see **Table S13**).

1. AcrAB-TolC upregulation. AcrAB-TolC is the paradigm efflux pump of the RND family of multidrug efflux pumps [24]. AcrB is the innermembrane structural component of the pump, while AcrR is a negative regulator of the AcrAB operon and PhoQ is a positive regulator of TolC, the outermembrane component of the pump [24]. We found a total of 111 SNPs and indels in *acrB*, *acrR* and *phoQ* in the 96 CTX-treated populations, and only one in the eight tetracycline-treated populations, suggesting that the pump has little affinity for tetracycline. Further, AcrB has a transmembrane and periplasmic domain and determines the pump's specificity [24]. Because only one of the 61 mutations affecting this gene inactivates the gene (one small population has a frameshift mutation), it is likely that these mutations either enhance the affinity for CTX. Mutations in *acrR* include several frameshift and nonsense mutations, consistent with its role in repressing expression of *acrAB*, while all eight mutations in *phoQ* are nonsynonymous SNPs, which presumably enhance expression of TolC [24]. Therefore, mutations in *acrR*, *acrB* and *phoQ* all seem to be involved in the upregulation of AcrAB-TolC and increasing its affinity for CTX.
2. Downregulation of plasmid copy number. The copy number of plasmid pACTEM is low (~10 copies/cell) due to relaxed regulation of its p15A origin of replication (ori) [12]. Inactivating mutations in *pcnB* and *polA* have been found to decrease copy number of plasmids with related p15A ori [25]. Because the mutations in *pcnB* and *polA* likely inactivated both gene functions (they include frameshift mutations and stop codons), and since lowered copy number was expected to be beneficial given that TEM1 β-lactamase was expressed from a strong pTac promoter and derepressed with IPTG, we expected these mutations to have decreased plasmid copy number. To test this, we compared TEM expression levels of the evolved clones from the 24 large populations, in which the plasmid was replaced by the ancestral pACTEM1 plasmid, with those in the two ancestral strains, using nitrocefin assays (see SM section 3, Experimental methods and results: "TEM β-lactamase activity assays"). The clone from population L18 was the only one without mutations in either *pcnB* or *polA*, and the clones from this population and population L5 (which had a 210-bp deletion in *pcnB*) were the only two showing lower β-lactamase expression than the ancestral strains (see **Fig. S4**). Therefore, the mutations inactivating *pcnB* and *polA* likely provided a fitness benefit by down-regulating plasmid copy number.
3. Transcription regulation. MarR is a DNA-binding transcriptional repressor involved in the regulation of many biological processes, including antibiotic resistance [26]. The 42 SNPs and indels affecting

*marR* include many nonsense and frame-shift mutations. Inactivation of MarR is expected to both upregulate AcrAB-TolC efflux and downregulate OmpF [24]. SlyA is a related transcription factor involved in the regulation of virulence, the silencing of horizontally acquired genes and repression of small-molecule efflux systems [27]. The mutations we observed inactivate SlyA and occur only in CTX-treated populations, suggesting that they may enhance CTX efflux. Non-synonymous SNPs in *rpoB* and *rpoD*, at different positions from those we found in our clones, have recently been reported to increase resistance to the cephalosporin ceftriaxole in *Neisseria ghonorrhoeae* [28]. DNA gyrase is not known for its role in β-lactam resistance. Mutations in *rpoABCD* and *gyrAB* were found only in the CTX-treated populations and did not include nonsense or frame-shift mutations. Moreover, two clones had the same Val771Gly mutation in *gyrA* and six clones had mutations at the same position in *rpoD* (of which four shared Asp445Glu), indicating selective benefits associated with changes in the regulation of gene expression.

4. OmpF downregulation. OmpF is the major outermembrane porin in *E. coli* B, and involved in the influx of β-lactams and other antibiotics [29]. The fact that *ompF* is deleted also in seven of the eight control populations challenged with tetracycline only, suggests tetracycline also enters through this porin. EnvZ-OmpR is a two-component regulatory system involved in the response to changes in osmolarity and positively regulating the expression of OmpF [29]. Since the mutations affecting all three genes include inactivating mutations, such as frameshift and nonsense mutations, mutations affecting *ompF*, *ompR* and *envZ* likely downregulate or abolish expression of OmpF.
5. Target alteration. Penicillin-binding protein 3 (PBP3), encoded by *ftsI*, is a transpeptidase catalyzing the final step of cell-wall biosynthesis by cross-linking peptidoglycan strands. It is also the main target to which CTX binds [30], which results in the inhibition of cell-wall synthesis and further toxic downstream effects [31]. Of the 58 mutations affecting *ftsI*, 56 were nonsynonymous SNPs and two in-frame (3-bp) indels. Moreover, we found high repeatability of mutations at the nucleotide level, with mutations at nine amino-acid positions occurring in more than one population, including 12 at position 536 (all Glu536Leu) and nine at position 167 (all Arg167Cys). At positions 311 (4 clones) and 545 (1 clone) mutations were also found in a previous study using CTX selection [30]. These PBP3 mutations, therefore, provide clear selective benefits, most likely by reducing CTX binding, while in theory -- since PBP's are evolutionary related to β-lactamases – some of these mutations might enhance the hydrolysis of CTX [30].
6. TEM deletion. All but two of the 45 deletions affecting TEM1 β-lactamase involved the 2,374-bp full deletion of $bla_{TEM1}$ and its repressor *lacI* from the plasmid (**Fig. 2C**); in two small populations, smaller deletions partially removing $bla_{TEM1}$ and its promoter were observed. All deletions are thus expected to fully cancel expression of TEM1. The $bla_{TEM1}$ and *lacI* loci are deleted by a single recombination event mediated by two homologous 185-bp sequences up and downstream of the two adjacent genes in the pACTEM1 plasmid. The deletion of these two genes is not driven by a selective benefit (**Fig. S3**), but by a high deletion rate and genetic hitchhiking with beneficial mutations. Consistent with this scenario, in the five populations with time-resolved information where the TEM deletion

occurs (populations S1, S14, L2, L3 and L5, **Fig. 4A**), it always spreads together with other mutations.

7. <u>TEM activation</u>. Besides the common deletion of *bla*$_{TEM1}$ and *lacI* and six *bla*$_{TEM1}$ promoter mutations (see above), 32 of the remaining 33 other mutations affecting TEM1 β-lactamase were nonsynonymous SNPs (the remaining mutation was a frameshift indel in a small population). The 32 SNPs are all known from clinical and/or laboratory studies [32], and show remarkable convergence, with 28 shared by multiple clones. These include known largest-benefit mutation Gly238Ser [33], shared by clones from 13 populations, and mutations at amino-acid positions 104, 164, 240, 241 and 265, shared by at least two clones. Eight large and two small populations had at least two SNPs in *bla*$_{TEM}$, including known adaptive combinations of Gly238Ser with Glu104Lys (3x) and Thr265Met (2x), and Arg164Ser with Ala237Thr [34]. Clearly, these mutations provided substantial fitness benefits by enhancing TEM's activity against CTX (see e.g. **Fig. S3**).
8. <u>Upregulation of outer-membrane vesicles</u>. NlpI is an outer-membrane-anchored lipoprotein, whose inactivation is known to enhance β-lactam resistance and increase the production of outer-membrane vesicles (OMVs) [35], as well as upregulate PBP4 during log phase and Spr during stationary phase [36]. Increased production of OMVs is thought to protect against stressors targeting the outer membrane via absorption [37]. Recently, the protective effect of OMVs against β-lactams such as CTX was shown to depend on the expression of an active β-lactamase [38]. Consistent with this scenario, the mutations we observed inactivated NlpI and were positively associated with TEM-activating SNPs (**Fig. S19**). Therefore, we expect that the benefit of *nlpI*-inactivating mutations in our populations resulted from the more effective removal of CTX via the increased production of OMVs.
9. <u>TEM expression</u>. The six mutations occurring upstream of *bla*$_{TEM1}$ occur all in the Tac promoter region (80-11bp upstream of the transcription start site), ranging from position -59 to -19, and including two at the same position (-41). It is unclear whether they increased or decreased expression of TEM β-lactamase. The strain we used has a copy of *lacI*, the gene encoding the LacI repressor of *bla*$_{TEM}$, both in the chromosome and on plasmid pACTEM. In total 16 SNPs and 25 large chromosomal deletions affect at least one *lacI* copy across the 96 populations (in addition to 45 combined deletions of *bla*$_{TEM1}$ and *lacI* from the plasmid). The deletion varied in size from 17,365-55,080 bp and affected at least 14 other genes (**Table S4**) and also occurred in one of the control populations that lost the pACTEM plasmid. Moreover, no frameshift or nonsense mutations affecting *lacI* were observed (which would upregulate TEM expression), and at three amino-acid positions mutations were observed in two populations. Furthermore, we know that a majority of populations have mutations in *pcnB* or *polA* that lower TEM expression (**Fig. S4**), presumably by lowering plasmid copy number, suggesting lower TEM expression is adaptive. Therefore, we expect that the *lacI* mutations downregulate TEM expression, either through weaker binding of LacI to lactose analog IPTG or tighter binding to the *bla*$_{TEM1}$ promoter. This would imply that the large deletions removing *lacI* may not provide a fitness benefit, at least not due to the removal of *lacI*.

We then looked for associations among mutations affecting these nine functional targets, in order to reveal possible adaptive trajectories in our populations, following the approach of Tenaillon et al. [19]. We excluded SVs from this analysis, because their adaptive role is unclear. However, we did include the deletion of *TEM*, as this particular SV is a highly repeated event with clear adaptive consequences (**Fig. S20**). We looked for interactions between functional targets using two approaches: (1) a Spearman rank correlation ($\rho$) between all functional units, and (2) Lewontin's normalized coefficient of linkage disequilibrium ($D'$) [37]. Whereas different mutational trajectories are evident for the large populations, for the small populations this is not the case (**Fig. S19**).

**Table S13**. Functional targets based on mutations affecting multiple-hit genes in the 96 CTX-treated populations.

| Functional target | Gene | SNP[1] | | | Indel/IS insertion | | SV | | **Total number** |
|---|---|---|---|---|---|---|---|---|---|
| | | Syn | NonSyn[2] | Reg | Coding | Reg | Deletion (>1kbp) | Duplication (>1kbp) | |
| 1. AcrAB-TolC upregulation | *acrB* | 0/0 | 43/16 | 0/0 | 1/1 | 0/0 | 0/0 | 6/0 | **50/17** |
| | *acrR* | 0/0 | 17(3*)/7(1*) | 2/0 | 13/3 | 0/0 | 1/0 | 4/0 | **37/10** |
| | *phoQ* | 0/0 | 4/4 | 0/0 | 0/0 | 0/0 | 0/0 | 0/0 | **4/4** |
| 2. Downregulation plasmid copy number | *pcnB* | 1/0 | 52(9*)/14(1*) | 0/2 | 14/6 | 2/0 | 0/0 | 0/0 | **69/22** |
| | *polA* | 0/0 | 5/2 | 0/0 | 1/0 | 0/0 | 0/0 | 15/2 | **21/4** |
| 3. Transcription regulation | *marR* | 1/0 | 15(2*)/4 | 2/0 | 3/0 | 15/2 | 0/0 | 1/0 | **37/6** |
| | *slyA* | 0/0 | 8(1*)/1(1*) | 0/0 | 5/0 | 2/0 | 0/0 | 0/0 | **15/1** |
| | *gyrAB* | 0/1 | 12/1 | 0/0 | 1[5]//0 | 0/0 | 0/0 | 2/0 | **15/2** |
| | *rpoABCD* | 0/0 | 15/7 | 0/0 | 0/0 | 0/0 | 0/0 | 0/0 | **15/7** |
| 4. OmpF downregulation | *ompR* | 0/0 | 19(3*)/4 | 0/0 | 9/1 | 1/2 | 1/0 | 1/0 | **31/7** |
| | *ompF* | 0/0 | 8(2*)/0 | 3/0 | 4/1 | 4/2 | 0/0 | 0/0 | **19/3** |
| | *envZ* | 0/0 | 9(1*)/8(4*) | 0/0 | 2/4 | 0/0 | 2/0 | 0/0 | **13/12** |
| 5. Target alteration | *ftsI* | 0/0 | 23/33 | 0/0 | 2[6]/0 | 0/0 | 0/0 | 0/0 | **25/33** |
| 6. TEM deletion[3] | *lacI* + *bla*$_{TEM}$ | | | | | | 39/6 | | **39/6** |
| 7. TEM activation[3] | *bla*$_{TEM}$ | | 7/25 | | | | | | **7/25** |
| 8. Upreg of outer-membrane vesicles | *nlpI* | 0/0 | 3(2*)/4(1*) | 0/0 | 9/9 | 0/1 | 0/0 | 0/0 | **12/14** |
| 9. TEM expression[3] | *lacI* | 0/0 | 8/8 | 0/0 | 0/0 | 0/0 | 21/4[4] | 0/0 | **29/12** |
| | *bla*$_{TEM}$ | 0/0 | | 0/6 | 0/0 | 1/0 | | 0/0 | **1/6** |
| **Total** | | **2/1** | **248(23*)/138(8*)** | **7/8** | **64/25** | **25/7** | **64/10** | **29/2** | **439/191** |

[1] Mutation numbers for Small/Large populations, respectively. Syn: synonymous, NonSyn: nonsynonymous, Reg: mutation in putative regulatory sequence of gene

(2) (*) indicate nonsense mutations.

(3) For the *lacI* and $bla_{TEM}$ genes, each mutation class is assigned to a different functional target. Grey shading indicates mutation classes that are not compatible with the functional target.

(4) Deletions of chromosomal *lacI* copy.

(5) An in-frame 6-bp insertion.

(6) Both indels are 3-bp in-frame deletions.

#### *Analysis of evolutionary trajectories*

To identify evolutionary trajectories we looked for associations between the mutations assigned to these nine functional targets, following the approach of Tenaillon et al. [19]. We were particularly interested in the effects of the TEM deletion and TEM activation on the trajectory followed. There were no clear associations for small populations, whereas large populations revealed two alternative trajectories (**Fig. S19**) which had adaptive consequences, affecting the resistance levels (**Fig. S20**).

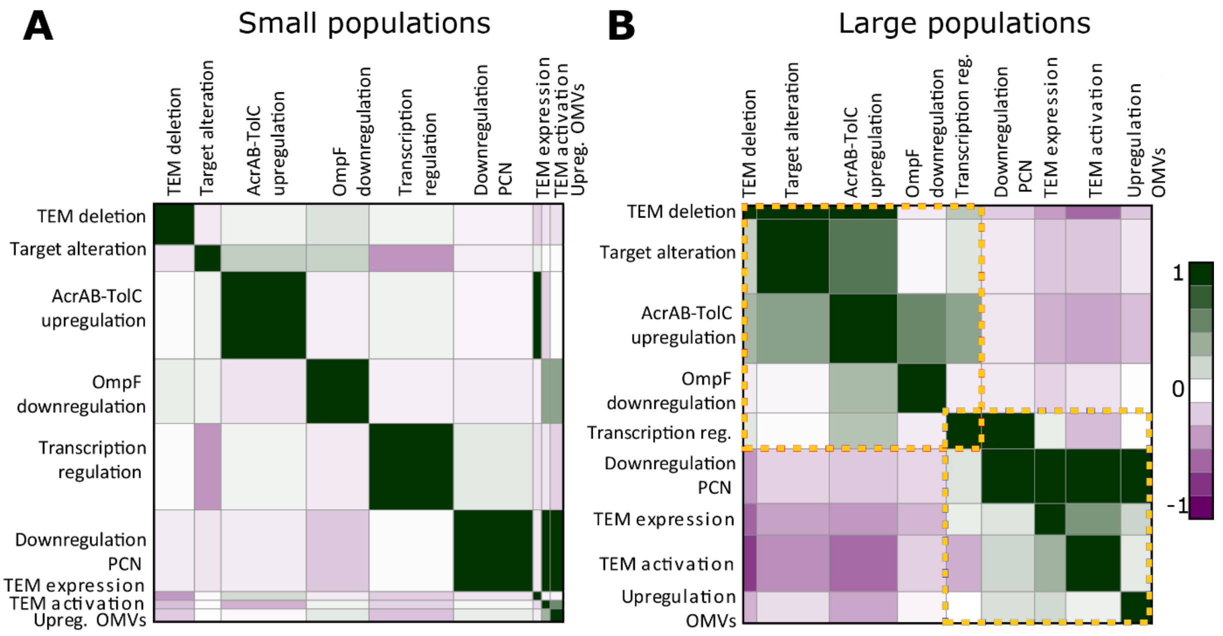


**Fig. S19:** Associations between functional targets with frequent mutations (see **Fig. 3C** and **Table S13**) in small (panel A) and large (panel B) populations, based on Spearman correlation (above diagonal) and normalized linkage disequilibrium [39] (below diagonal). Height and width of boxes reflects mutation frequencies, colors indicate negative (purple) to positive (green) associations. The yellow dashed lines in the panel for large populations highlight two alternative trajectories involving the deletion or activation of TEM1 β-lactamase.

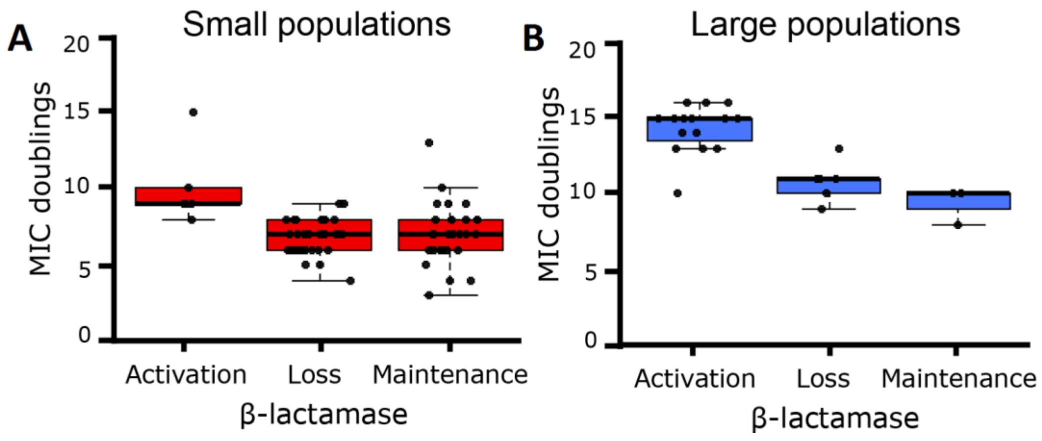


**Fig. S20:** MIC doublings of analyzed evolved clones for genotypes that activate, delete or maintain TEM1, for small (panel A) and large (panel B) populations. TEM1 maintenance indicates that TEM1 was neither activated or deleted.

### References


1. Gerrish, P.J. and R.E. Lenski, *The fate of competing beneficial mutations in an asexual population.* Genetica, 1998. **102/103**: p. 127-144.
2. Good, B.H., et al., *Distribution of fixed beneficial mutations and the rate of adaptation in asexual populations.* Proceedings of the National Academy of Sciences USA, 2012. **109**: p. 4950–4955.
3. Park, S.-C., D. Simon, and J. Krug, *The speed of evolution in large asexual populations.* Journal of Statistical Physics, 2010. **138**: p. 381-410.
4. Good, B.H. and M.M. Desai, *Deleterious Passengers in Adapting Populations.* Genetics, 2014. **198**(3): p. 1183-1208.
5. Schiffels, S., et al., *Emergent Neutrality in Adaptive Asexual Evolution.* Genetics, 2011. **189**: p. 1361–1375.
6. McCandlish, D.M. and A. Stoltzfus, *Modeling Evolution Using the Probability of Fixation: History and Implications.* The Quarterly Review of Biology, 2014. **89**(3): p. 225-252.
7. Yampolsky, L.Y. and A. Stoltzfus, *Bias in the introduction of variation as an orienting factor in evolution.* Evolution & Development, 2001. **3**(2): p. 73-83.
8. Jain, K., J. Krug, and S.-C. Park, *Evolutionary advantage of small populations on complex fitness landscapes.* Evolution, 2011. **65**: p. 1945–1955.
9. Svensson, E.I. and D. Berger, *The Role of Mutation Bias in Adaptive Evolution.* Trends in Ecology & Evolution, 2019. **34**(5): p. 422-434.
10. Gomez, K., J. Bertram, and J. Masel, *Mutation bias can shape adaptation in large asexual populations experiencing clonal interference.* Proceedings of the Royal Society B: Biological Sciences, 2020. **287**(1937): p. 20201503.
11. Lenski, R.E., et al., *Long-term experimental evolution in Escherichia coli. I. Adaptation and divergence during 2,000 generations.* American Naturalist, 1991. **138**: p. 1315-1341.
12. Barlow, M. and B.G. Hall, *Predicting evolutionary potential: In vitro evolution accurately reproduces natural evolution of the TEM beta-lactamase.* Genetics, 2002. **160**: p. 823-832.
13. de Visser, J.A.G.M., et al., *Diminishing returns from mutation supply rate in asexual populations.* Science, 1999. **283**: p. 404-406.
14. Martin, M., *Cutadapt removes adapter sequences from high-throughput sequencing reads.* EMBnet.journal, 2011. **17**(1): p. 10-11.
15. Bolger, A.M., M. Lohse, and B. Usadel, *Trimmomatic: A flexible trimmer for Illumina Sequence Data.* Bioinformatics, 2014. **30**(15): p. 2114-2120.
16. Langmead, B. and S.L. Salzberg, *Fast gapped-read alignment with Bowtie 2.* Nature Methods,

2012. **9**(4): p. 357-359.
17. Li, H., et al., *The Sequence Alignment/Map format and SAMtools.* Bioinformatics, 2009. **25**(16): p. 2078-2079.
18. Lee, H., et al., *Rate and molecular spectrum of spontaneous mutations in the bacterium Escherichia coli as determined by whole-genome sequencing.* Proceedings of the National Academy of Sciences, 2012. **109**: p. E2774–E2783.
19. Tenaillon, O., et al., *The Molecular Diversity of Adaptive Convergence.* Science, 2012. **335**: p. 457-461.
20. Miller, C.A., et al., *Visualizing tumor evolution with the fishplot package for R.* BMC Genomics, 2016. **17**(1): p. 880.
21. Nowak, S., et al., *Multidimensional Epistasis and the Transitory Advantage of Sex.* PLoS Comput Biol, 2014. **10**(9): p. e1003836.
22. Bishop, C.M., *Mixture density networks*. 1994, Birmingham: Aston University.
23. Bishop, C.M., *Pattern Recognition and Machine Learning*. 2011, New York: Springer.
24. Du, D., et al., *Multidrug efflux pumps: structure, function and regulation.* Nature Reviews Microbiology, 2018. **16**(9): p. 523-539.
25. Chakravartty, V. and J.E. Cronan, *A series of medium and high copy number arabinose-inducible Escherichia coli expression vectors compatible with pBR322 and pACYC184.* Plasmid, 2015. **81**: p. 21-26.
26. Ellison, D.W. and V.L. Miller, *Regulation of virulence by members of the MarR/SlyA family.* Current Opinion in Microbiology, 2006. **9**(2): p. 153-159.
27. Will, W.R., et al., *The Evolution of SlyA/RovA Transcription Factors from Repressors to Countersilencers in <em>Enterobacteriaceae</em>.* mBio, 2019. **10**(2): p. e00009-19.
28. Palace, S.G., et al., *RNA polymerase mutations cause cephalosporin resistance in clinical Neisseria gonorrhoeae isolates.* eLife, 2020. **9**: p. e51407.
29. Choi, U. and C.-R. Lee, *Distinct Roles of Outer Membrane Porins in Antibiotic Resistance and Membrane Integrity in Escherichia coli.* Frontiers in microbiology, 2019. **10**: p. 953-953.
30. Sun, S., M. Selmer, and D.I. Andersson, *Resistance to b-Lactam Antibiotics Conferred by Point Mutations in Penicillin-Binding Proteins PBP3, PBP4 and PBP6 in Salmonella enterica.* PLoS One, 2014. **9**: p. e97202.
31. Cho, H., T. Uehara, and Thomas G. Bernhardt, *Beta-Lactam Antibiotics Induce a Lethal Malfunctioning of the Bacterial Cell Wall Synthesis Machinery.* Cell, 2014. **159**(6): p. 1300-1311.
32. Salverda, M.L.M., J.A.G.M. de Visser, and M. Barlow, *Natural evolution of TEM-1 beta-lactamase: experimental reconstruction and clinical relevance* FEMS Microbiology Reviews, 2010. **34**: p. 1015–1036.
33. Schenk, M.F., et al., *Quantifying the adaptive potential of an antibiotic resistance enzyme.* PLoS Genetics, 2012. **8**: p. e1002783.
34. Salverda, M.L.M., et al., *Initial mutations direct alternative pathways of protein evolution.* PLoS Genetics, 2011. **7**(3): p. e1001321.
35. Kim, S.W., et al., *Outer membrane vesicles from β-lactam-resistant Escherichia coli enable the survival of β-lactam-susceptible E. coli in the presence of β-lactam antibiotics.* Scientific Reports, 2018. **8**(1): p. 5402.
36. Schwechheimer, C., D.L. Rodriguez, and M.J. Kuehn, *NlpI-mediated modulation of outer membrane vesicle production through peptidoglycan dynamics in Escherichia coli.* MicrobiologyOpen, 2015. **4**(3): p. 375-389.
37. Manning, A.J. and M.J. Kuehn, *Contribution of bacterial outer membrane vesicles to innate bacterial defense.* BMC Microbiology, 2011. **11**(1): p. 258.
38. Kim, S.W., et al., *The Importance of Porins and β-Lactamase in Outer Membrane Vesicles on the Hydrolysis of β-Lactam Antibiotics.* International Journal of Molecular Sciences, 2020. **21**(8): p. 2822.
39. Lewontin, R.C., *The interaction of selection and linkage. I. General considerations; heterotic models.* Genetics, 1964. **49**: p. 49-67.
